# NanopoReaTA: a user-friendly tool for nanopore-seq real-time transcriptional analysis

**DOI:** 10.1101/2022.12.13.520220

**Authors:** Anna Wierczeiko, Stefan Pastore, Stefan Mündnich, Anne M. Busch, Vincent Dietrich, Mark Helm, Tamer Butto, Susanne Gerber

## Abstract

**Summary:** Oxford Nanopore Technologies’ (ONT) sequencing platform offers an excellent opportunity to perform real-time analysis during sequencing. This feature allows for early insights into experimental data and accelerates a potential decision-making process for further analysis, which can be particularly relevant in the clinical context. Although some tools for the real-time analysis of DNA-sequencing data already exist, there is currently no application available for differential transcriptome data analysis designed for scientists or physicians with limited bioinformatics knowledge. Here we introduce NanopoReaTA, a user-friendly real-time analysis toolbox for RNA sequencing data from ONT. Sequencing results from a running or finished experiment are processed through an R Shiny-based graphical user interface (GUI) with an integrated *Nextflow* pipeline for whole transcriptome or gene-specific analyses. NanopoReaTA provides visual snapshots of a sequencing run in progress, thus enabling interactive sequencing and rapid decision-making that could also be applied to clinical cases.

**Availability:** https://github.com/AnWiercze/NanopoReaTA

## 1 Introduction

In standard sequencing experiments, practical steps and data analysis are usually performed independently, with the latter initiated by bioinformatics experts once sequencing is complete. Nowadays, new technologies such as Oxford Nanopore Technologies (ONT) offer a unique opportunity to start downstream analysis while sequencing is still ongoing (Amarasinghe et al., 2020, Wang et al., 2021). Some platforms, such as EPI2ME from ONT (https://labs.epi2me.io/) or minoTour (https://github.com/minoTour/minoTour, Munro et al. 2022), already provide real-time pipelines for rapid ONT data acquisition integrated into a user interface (UI), and thus accessible to users with limited bioinformatics skills. However, as these platforms ‘ focus mainly lies on the analysis of DNA sequencing data, there is a lack of real-time applications in the field of transcriptomics. Here we introduce NanopoReaTA, an on-demand toolbox for real-time transcriptomic analysis that provides rapid insight on RNA sequencing data from ONT. Users receive transcriptome-wide and gene-specific information directly while sequencing is still running, such as differences between conditions or expression levels of individual genes. In addition, implemented quality control features allow the user to monitor data variability during the ongoing sequencing process. Ultimately, the tool can provide frequent biologically relevant snapshots of the current sequencing run, which in turn can enable interactive fine-tuning of the sequencing run itself, facilitate decisions to abort the ongoing run to save time and material, e.g., when sufficient accuracy is achieved, or even accelerate the resolution of clinical cases with high urgency.

## 2 Material and methods

### 2.1 Test Data

NanopoReaTA has been tested on self-generated direct cDNA sequencing data from Hek293 and HeLa cells (Supplementary Information, Table S1 and Figure S1-S9).

### 2.2 Usage

NanopoReaTA can be launched directly after starting a sequencing run of cDNA or direct RNA via ONT ‘s sequencing software MinKNOW (Supplementary Information, Figure 1A and Figure S1). Within NanopoReaTA ‘s UI, the user will be guided through several configuration settings to extract all information required for data processing such as reference sequences, annotation files, output directory defined in MinKNOW (into which sequencing output is written), and more (Figure 1B, Figure S2A-C, Supplementary Information). Preprocessing of basecalled reads from a running or completed experiment is integrated into a *Nextflow* pipeline and can be started via a one-button-click within the UI (Figure 1C, Figure S2D, Di Tommaso et al. 2017). As soon as sequencing data are generated, the *Nextflow* pipeline automatically updates generated files, including gene counts or mapping files. Based on the output files from preprocessing, downstream analyses can be performed within the following tabs integrated into NanopoReaTA: “Overview “, “Gene-wise Analysis “ and “Differential Expression Analysis “ (Figure S4-6). The resulting figures can be constantly updated during sequencing (Figure S7-8). See more details in the Supplementary Information.

**Figure 1.**
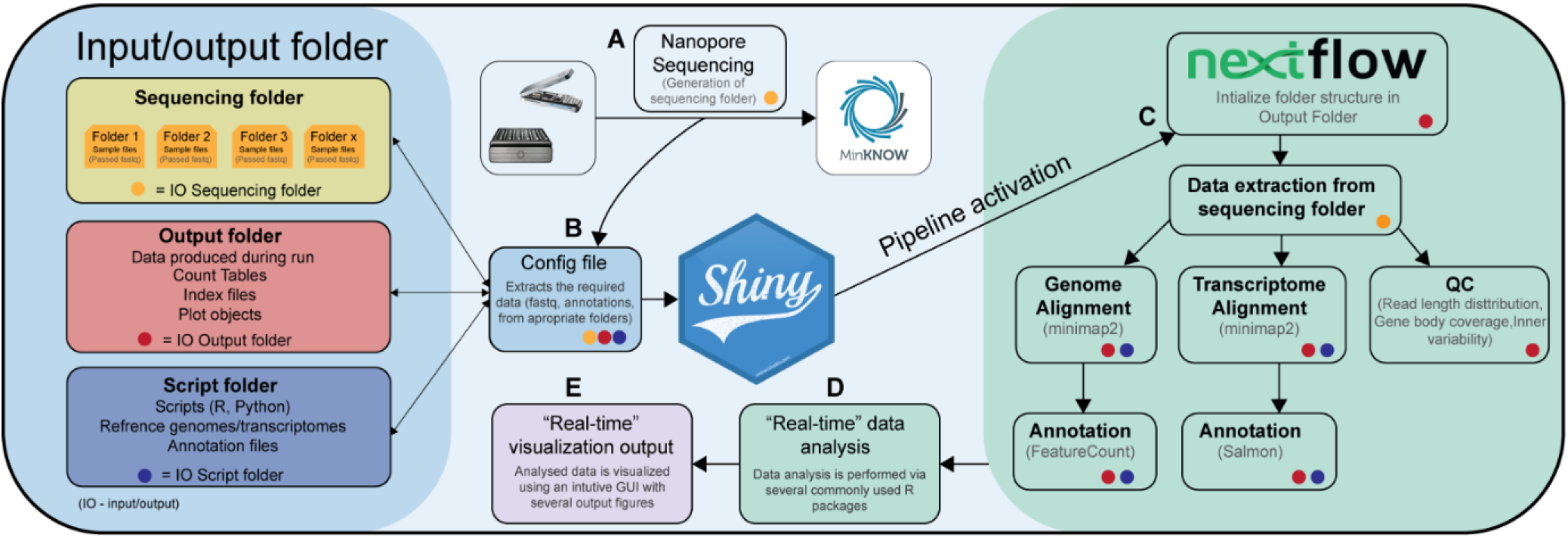
NanopoReaTA workflow. visualized as a graphical sketch. Detailed information on the individual modules are given in the Supplementary Information, as well as in the user manual on the GitHub repository: https://github.com/AnWiercze/NanopoReaTA

### 2.3 Preprocessing via *Nextflow*

The *Nextflow* pipeline takes all fastq files that pass the quality threshold defined in MinKNOW and performs genome and transcriptome alignment using minimap2 (Li, 2018) as well as feature quantification using FeatureCounts (Liao et al., 2014) and Salmon (Patro et al., 2017). In addition, we incorporated a quality control utility extracting sample- and group-wise read length distribution, variability measurements, genome/transcriptome coverage based on RSeQC (Wang et al., 2012), and gene count per iteration, enabling the assessment of specific quality metrics over time (Figure S7). See more details in the Supplementary Information.

### 2.4 Downstream analyses based on R

The subsequent downstream analyses are based on commonly used R packages such as DESeq2 (Love et al., 2014) for principal component analysis (PCA) and differential expression analysis of gene and transcript expression, and DEXSeq (Anders et al., 2012) and DRIMSeq (Robinson, & Nowicka 2016) for differential transcript usage (Figure 1D-E). In addition, gene body coverage and counts per sample and condition can be visualized for a subset of genes of interest (Figure 1E). All tables and figures can be downloaded via button clicks (Figure S3-6). See more details in the Supplementary Information.

### 2.5 Installation and requirements

NanopoReaTA can be installed on Linux and Windows via docker by pulling a pre-build docker image containing all package requirements. For installation, requirements, and user manual, please visit https://github.com/AnWiercze/NanopoReaTA. <u>*Hardware:*</u> 64GB RAM, 16 threads <u>*Software:*</u> Docker

## 3 Discussion

NanopoReaTA represents a real-time analysis toolbox that allows users to perform interactive transcriptional analyses of cDNA and direct RNA sequencing data in real-time via a user-friendly and intuitive user interface based on R Shiny. We aim to provide a tool that supports users from biological research and clinical diagnostics of transcriptomics by accelerating decision-making processes of future experiments or patient treatment, especially when time and money are limiting factors. For future perspectives, we envision that additional functions such as novel transcript detection, RNA modification detection, and integration of multi-omics levels in real-time can be integrated. NanopoReaTA is open source to also enable the scientific community to contribute such enhancements.

## Supporting information

NanopoReaTA - Proof of concept

NanopoReaTA - Final Results

## Author contributions

TB and SG conceived and supervised the project. AW and SP designed, implemented, and tested the GUI. AW, SP, VM and AB implemented GUI updates. SM performed the RNA isolation from Hek293 and HeLa and TB performed the direct cDNA library preparation. TB, AW, and SP wrote the manuscript. SG, MH, and SM edited the manuscript and provided valuable input and feedback in various discussions. All authors read and approved the final manuscript.

## References

Anders, S., Reyes, A., & Huber, W. (2012). Detecting differential usage of exons from RNA-seq data. Genome Research, 22(10), 2008–2017. https://doi.org/10.1101/GR.133744.111

Di Tommaso, P., Chatzou, M., Floden, E. W., Barja, P. P., Pa-lumbo, E., & Notredame, C. (2017). Nextflow enables re-producible computational workflows. Nature biotechnology, 35(4), 316–319. doi: 10.1038/nbt.3820.

Li, H. (2018). Minimap2: pairwise alignment for nucleotide sequences. Bioinformatics, 34(18), 3094–3100. https://doi.org/10.1093/BIOINFORMATICS/BTY191

Liao, Y., Smyth, G. K., & Shi, W. (2014). featureCounts: an efficient general purpose program for assigning sequence reads to genomic features. Bioinformatics, 30(7), 923–930. https://doi.org/10.1093/BIOINFORMATICS/BTT656

Love, M. I., Huber, W., & Anders, S. (2014). Moderated estimation of fold change and dispersion for RNA-seq data with DESeq2. Genome Biology, 15(12), 1–21. https://doi.org/10.1186/S13059-014-0550-8/FIGURES/9

Munro, R., Santos, R., Payne, A., Forey, T., Osei, S., Holmes, N., & Loose, M. (2022). minoTour, real-time monitoring and analysis for nanopore sequencers. Bioinformatics, 38(4), 1133–1135. https://doi.org/10.1093/BIOINFORMATICS/BTAB780

Robinson, M. D., & Nowicka, M. (2016). DRIMSeq: A Dirichlet-multinomial framework for multivariate count outcomes in genomics. F1000Research, 5. https://doi.org/10.12688/F1000RESEARCH.8900.2/DOI

Patro, R., Duggal, G., Love, M. I., Irizarry, R. A., & Kingsford, C. (2017). Salmon provides fast and bias-aware quantification of transcript expression. Nature Methods 2017 14:4, 14(4), 417–419. https://doi.org/10.1038/nmeth.4197

Wang, L., Wang, S., & Li, W. (2012). RSeQC: quality control of RNA-seq experiments. Bioinformatics, 28(16), 2184–2185. https://doi.org/10.1093/BIOINFORMATICS/BTS356

