## Supplementary Information for "NanopoReaTA: a user-friendly tool for nanopore-seq real-time transcriptional analysis"

**Materials and methods**

***Cell culture.*** HEK293 and HeLa cells were cultured and maintained in an incubator at 37°C and 5% CO2. HEK293 and HeLa cells were cultured in DMEM supplemented with 10% FBS, 1% penicillin-streptomycin, and 1% L-glutamine. Once the cells were confluent, the medium was removed and cells were washed once with 1 mL DPBS. The cells were resuspended with 0.5 mL Trizol and collected for Trizol RNA isolation.

***RNA isolation.*** For RNA isolation, following 5 min incubation in RT, 100 µL of chloroform was added. Samples were vortexed and incubated 2 minutes in RT. Sample were centrifuged 13000xg, 4 °C for 10 min and the upper aqueous phase was transferred into a new tube. Next, 250 µL of isopropanol was added and incubated for 15 min in RT for RNA precipitation. The samples were then centrifuged 13000 – 15000xg at 4 °C for 30 min and supernatant was discarded. RNA pellet was washed with cold 75% cold EtOH (stored at -20 °C) and centrifuged again 13000 – 15000xg at 4 °C for 30 min. Supernatant was discarded and the pellet was air dried. RNA pellet was resuspended with nuclease-free water and concentration was measured using nanodrop.

***Direct cDNA-native barcoding Nanopore Library preparation.*** Libraries were prepared according to Direct cDNA sequencing - native barcoding (SQK-DCS109 with EXP-NBD104). 2µg of total RNA were combined with 2.5µL of VNP, 10mM dNTPs and RNAse-free water to a final volume of 11 µL. Samples were Incubated at 65°C for 5 minutes and then snapped cool on a pre-chilled freezer block for 1 minute. Strand-switching buffer (5x RT Buffer µL, RNaseOUT 1 µL, Nuclease-free water 1µL, Strand-Switching Primer (SSP) 2 µL, final volume 8 µL) was added to each of snap-cooled samples and incubated at 42°C for 2 minutes. 1 µl of Maxima H Minus Reverse Transcriptase (ThermoFisher, cat # EP0751) was added to each sample (to a final volume of 20 µL) and incubated in thermocycler at 42°C for 90 min following by 85°C heat inactivation for 5 min. For RNA degradation, 1 µl RNase Cocktail Enzyme Mix (ThermoFisher, cat # AM2286) was added to each sample and incubated for 10 minutes at 37° C in a thermal cycler. The samples were cleaned up using 0.8× AMPure XP Beads (Agencourt, A63881) and eluted in 20 µL nuclease-free water. For second strand synthesis , 25 µL 2x LongAmp Taq Master Mix, 2 µL of PR2 Primer (10 μM) and 3 µL of nuclease-free water were added to reversed transcribed sample to final volume of 50 µL. The samples were incubated at 94 °C for 1 mins, 50 °C for 1 min and 65 °C for 15 min. The samples were cleaned up using 0.8× AMPure XP Beads (Agencourt, A63881) and eluted in 21 µL nuclease-free water. Following strand-switched cDNA quantification, samples were end-prepped using NEBNext Ultra II End Repair / dA-tailing Module NEB, cat # E7546) (cDNA sample 20 µL, Nuclease-free water 30 µL, Ultra II End-prep reaction buffer 7 µL, Ultra II End-prep enzyme mix 3 µL, final volume 60 µL) and incubated at 20°C for 10 minutes and 65°C for 10 minutes. The samples were cleaned up using 1× AMPure XP Beads and eluted in 22.5 µL nuclease-free water. End-prepped DNA were barcoded using Native Barcoding Expansion 1-12 (EXP-NBD104, ONT)( End-prepped DNA 22.5 µL, Native Barcode 2.5 µL, Blunt/TA Ligase Master Mix 25 µL, final volume 50 µL). The samples were cleaned up using 1× AMPure XP Beads and eluted in 26 µL nuclease-free water. For adapter ligation, barcoded samples were pooled (total 65 µL) and incubated with 5 µL Adapter Mix II (AMII), 20 µL NEBNext Quick Ligation Reaction Buffer (5X) and 10 µL Quick T4 DNA Ligase (NEB, cat # E6056) toa final volume of 100 µL. Lastly, samples were cleaned up using 0.5× AMPure XP Beads and washed twice with Wash Buffer (WSB, ONT) and eluted in 13 µL Elution Buffer (EB, ONT). Eluted sample were mixed with sequencing buffer and loading beads before loading onto a primed R9.4.1 flowcell and rum on MinION decive.

**Nextflow pipeline processing**

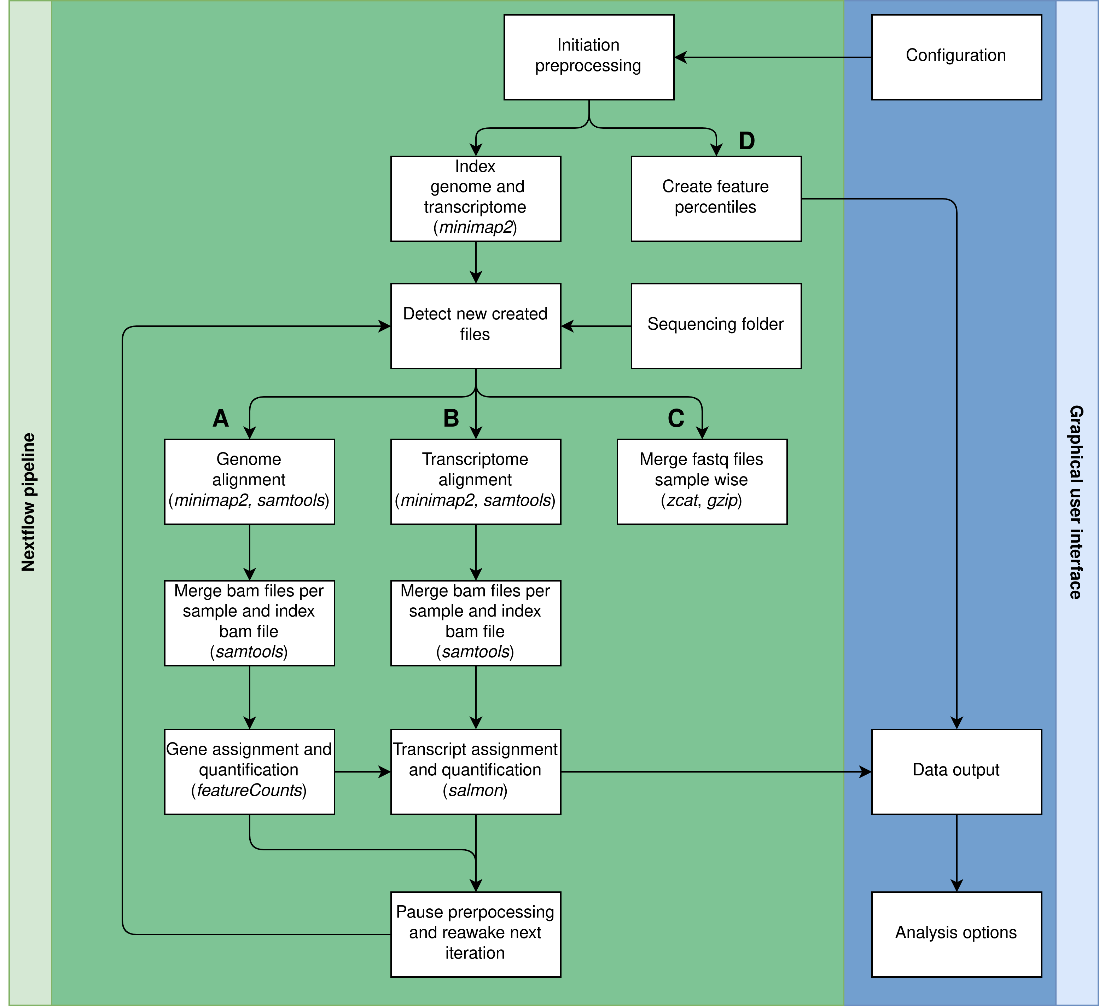
The *nextflow* (Di Tommaso 2017) pipeline is processing batches of fastq files in an iterative manner. By default, MinKNOW produces fastq files with 4000 reads. At the beginning of each iteration the *nextflow* pipeline checks if new files have been created in the determined output folders of MinKNOW. The pipeline processes a maximum of 30 files per iteration, which corresponds to 120.000 reads per iteration. The preprocessing pipeline runs with four parallel operations. All the operations have to finish in order to start a new preprocessing iteration. Each operation is programmed in an additive manner, which only works with newly created data and summarizes the outcome with the already processed data. The latter allows a time efficient performance. Data will be stored in a predefined run directory. In case of an unexpected interruption of the application, the pipeline cleans up corrupted files and recovers all the processed data. The preprocessing can be paused at the end of each iteration to allow a real-time analysis of meanwhile collected data.

**Supplementary Information Diagram 1. (A)** Genome alignment, annotation and quantification of gene expression. Indexing of genome alignment is performed a single time when the preprocessing is started. Genome alignment is performed with minimap2 (Li et al. 2018), BAM-files are sorted by using functionalities of samtools. Sorted BAM-files of each sample are then seperately merged via samtools merge and indexed via samtools index. The summarized BAM-files are used as input for featureCounts (Liao et al. 2014) for annotation and quantification of gene expression. The created count files are saved as semicolon separated CSV files. The count files of each sample are summarized to a countfile including each sample. Number of genes counted and the variability of gene counts is determined on basis of the constructed count files. **(B)** Transcriptome alignment, annotation and quantification of gene expression. Indexing of transcriptome is performed a single time when the preprocessing is started. Transcriptome alignment is performed with minimap2, BAM-files are sorted by using functionalities of samtools. Sorted BAM-files of each sample are then seperately merged via samtools merge. The merged bamfile becomes indexed. The summarized BAM-files are used as input for salmon for annotation and quantification of gene expression. The created count files are saved as semicolon separated CSV files. **(C)** Merge of single fastq files for each sample. Every detected fastq file will be merged to its respective sample fastq file using bash functionalities like cat/zcat and gzip. **(D)** During preparation for Gene Body Coverage, GeneBody pecentiles are determined at the first iteration of a newly startet preprocessing pipeline. This step only runs a single time at the beginning of the preprocessing pipeline.

Important commands

(A)

minimap2 --MD -ax splice (-uf –k14) -d MT-genome_ont.mmi genome_fasta

minimap2 --MD -ax splice (-uf –k14) -t ${params.threads} ${params.run_dir}MT-human_ont.mmi $i | samtools view -hbS -F 3844 | samtools sort > ${outname}

samtools merge all_gene.bam ${bam_files_to_merge} -f -h ${bam_files_to_merge} --threads ${params.threads} -c –p

samtools index alle_gene.bam

featureCounts -a ${genome_gtf} -F 'GTF' -L -T ${params.threads} -o merged_fc.csv all_gene.bam

python ${params.script_dir}merge_all_fc.py ${data_string} merged_all.csv

python ${params.script_dir}infer_experiment_absolute_gene_amount.py -s ${params.run_dir}merged_all.csv -m "" -o exp_genes_counted_per_sample.csv

python ${params.script_dir}infer_experiment_inner_variability.py -s ${params.run_dir}merged_all.csv -m "" -d ${params.run_dir}inner_variability_plot.csv -o }inner_variability_per_sample.csv

(B)

minimap2 --MD -ax map-ont -d MT-transcript_ont.mmi $params.transcriptome_fasta

minimap2 --MD -ax map-ont -t ${params.threads} ${params.run_dir}MT-human_transcript_ont.mmi ${params.run_dir} $i

samtools merge ${run_dir}/${sample}/salmon/all.bam ${sample}*.bam -f -h ${sample}*.bam --threads ${params.threads} -c -p

samtools index ${run_dir}/${sample}/all.bam

salmon quant -t ${params.transcriptome_fasta} -l A -a ${params.run_dir}/${sample}/salmon/all.bam -o ${params.run_dir}/${sample}/salmon/ -g ${params.genome_gtf} -p ${params.threads}

python ${params.script_dir}merge_all_salmon.py ${data_string} ${params.run_dir}salmon_merged_tpm.csv ${params.run_dir}salmon_merged_absolute.csv

(C)

zcat \${data_to_process} | gzip -c >> \${run_dir}\${par_basis}/merged_fastq/\${par_basis}.fastq.gz | \

cat \${data_to_process} >> \${run_dir}\${par_basis}/merged_fastq/\${par_basis}.fastq

(D)

python ${params.script_dir}createFeaturePercentiles.py --bed ${params.bed_file} --output_dir ${params.run_dir}

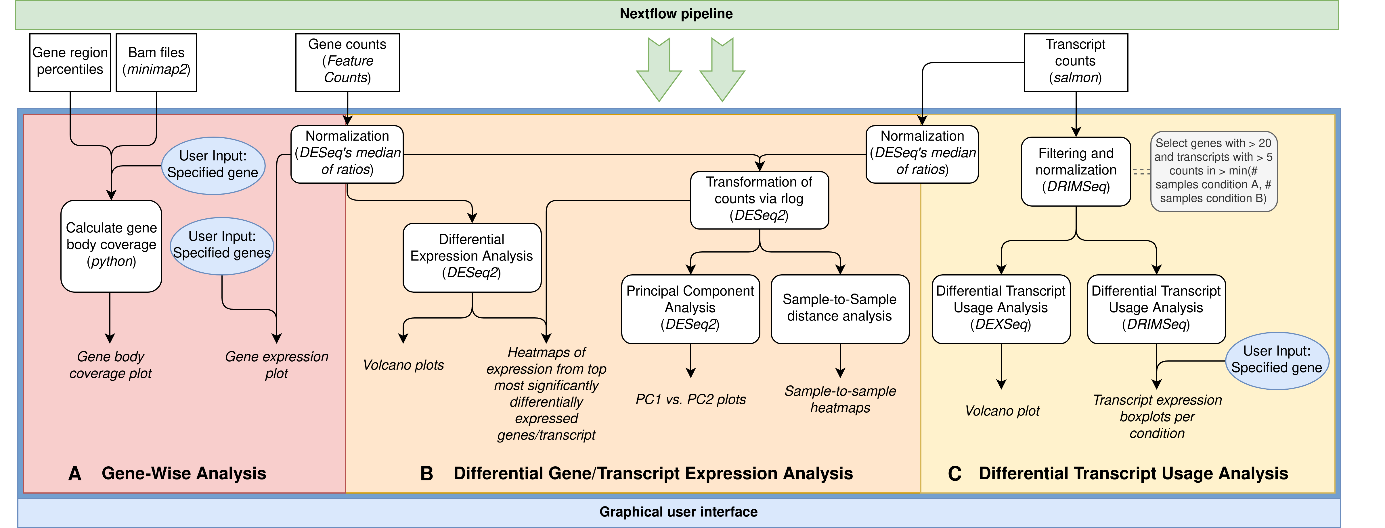
**Analysis in Rshiny**

**Supplementary Information Diagram 2.** Within the app three main analysis types can be applied on the output of the preprocessing pipeline by nextflow: **(A)** Gene-wise analysis: 1. The user can plot the gene-specific gene-body coverage utilizing the *gene region percentiles* and the mapping output - *bam files -* generated within the nextflow pipeline. The gene region is devided into 100 sub-regions/bins and the percentage of reads mapped onto each sub-region for individual samples or averaged over all samples from one condition is calculated and plotted. The approach for the gene body coverage analysis is based on the geneBodyCoverage.py function from RSeQC [Wang et al., 2012]. 2. Moreover, the gene expression/read counts for specified genes can be plotted as boxplots or barplots before and after normalization of *read counts* calculated by FeatureCounts [Liao et al. 2014]. **(B)** Differential gene/transcript Expression Analysis: Differential expression analysis of genes (DGE) and/or transcripts (DTE) can be executed. For gene level analysis, the raw gene counts from FeatureCounts and rounded estimated transcript counts from salmon are first normalized by DESeq2’s median-of-ratios approach and then tested for differential expression between the provided conditions using the Wald-Test provided by DESeq2. The order of testing (condition A vs. condition B or condition B vs. condition A) can be changed at the metadata settings page by changing the order of selection of conditions. This has an influence on the sign of the log2FoldChange values (log2(CondA/CondB) or log2(CondB/CondA)). In addition to the differenital expression analysis, a principal component analysis (PCA) and sample-to-sample euclidean distance analysis will be applied on normalized gene/transcript counts after transformation by the regularized logarithm (rlog) and visualized by plotting PC1 against PC2 and heatmap, respectively. Additionally, the rlog transformed counts are used to plot the expression of the top 20 most significant genes/transcripts by heatmap. **(C)** Furthermore, the user can make a differential transcript usage (DTU) analysis using DEXSeq [Anders et al., 2012] and DRIMSeq [Robinson & Nowicka 2016]. Here, we first filter and normalize the transcript counts from salmon [Patro et al. 2017] using DRIMSeq - transcript expression > 5 and gene expression > 20 reads in at least the number of samples of the smaller condition. The filtered and normalized transcript counts are then tested for differential transcript usage within each gene between the specified conditions by both DEXSeq and DRIMSeq. The transcript-based log2foldChange values from DEXSeq’s analysis will be plotted against the adjusted p-value (=volcano plot) to get a quick overview of the most significant differentially used transcripts with the highest effect size (log2FoldChange). From the DRIMSeq output, the user can plot the proportions of transcripts for a user-specified gene found in sample using boxplots per condition. Significantly differentially used transcripts are highlighted by a red line above the boxplots. The DTU workflow is based on the method article published by Love et al. in 2018.

References

Anders, S., Reyes, A., & Huber, W. (2012). Detecting differential usage of exons from RNA-seq data. *Genome Research*, *22*(10), 2008–2017. <https://doi.org/10.1101/GR.133744.111>

Di Tommaso, P., Chatzou, M., Floden, E. W., Barja, P. P., Pa-lumbo, E., & Notredame, C. (2017). Nextflow enables re-producible computational workflows. Nature biotechnology, 35(4), 316-319. doi: 10.1038/nbt.3820.

Li, H. (2018). Minimap2: pairwise alignment for nucleotide sequences. *Bioinformatics*, *34*(18), 3094–3100. https://doi.org/10.1093/BIOINFORMATICS/BTY191

Liao, Y., Smyth, G. K., & Shi, W. (2014). featureCounts: an efficient general purpose program for assigning sequence reads to genomic features. *Bioinformatics*, *30*(7), 923–930. https://doi.org/10.1093/BIOINFORMATICS/BTT656

Love, M. I., Huber, W., & Anders, S. (2014). Moderated estimation of fold change and dispersion for RNA-seq data with DESeq2. *Genome Biology*, *15*(12), 1–21. <https://doi.org/10.1186/S13059-014-0550-8/FIGURES/9>

Robinson, M. D., & Nowicka, M. (2016). DRIMSeq: A Dirichlet-multinomial framework for multivariate count outcomes in genomics. *F1000Research*, *5*. https://doi.org/10.12688/F1000RESEARCH.8900.2/DOI

Patro, R., Duggal, G., Love, M. I., Irizarry, R. A., & Kingsford, C. (2017). Salmon provides fast and bias-aware quantification of transcript expression. *Nature Methods 2017 14:4*, *14*(4), 417–419. https://doi.org/10.1038/nmeth.4197

Wang, L., Wang, S., & Li, W. (2012). RSeQC: quality control of RNA-seq experiments. *Bioinformatics*, *28*(16), 2184–2185. <https://doi.org/10.1093/BIOINFORMATICS/BTS356>
