## Supplementary Tables and Figures for "NanopoReaTA: a user-friendly tool for nanopore-seq real-time transcriptional analysis"

**Supplementary Table 1 - Data Used in the Study**

| **Sample alias** | **Cell Line** | **Barcodes** | **protocol** | **flowcell** | **kit** | **Platform** | **fastq files** |
| --- | --- | --- | --- | --- | --- | --- | --- |
| Hek293_directcDNA_replicate1_run1 | Hek293 | Barcode1 | direct-cDNA | FLO-MIN106D | SQK-DCS109 + EXP-NBD104 | MinION | FAQ51879_pass_barcode01_  03abc106_254e4132_*****.fastq.gz |
| Hek293_directcDNA_replicate2_run1 | Hek293 | Barcode2 | direct-cDNA | FLO-MIN106D | SQK-DCS109 + EXP-NBD104 | MinION | FAQ51879_pass_barcode02_  03abc106_254e4132_*****.fastq.gz |
| HeLa_directcDNA_replicate1_run1 | HeLa | Barcode3 | direct-cDNA | FLO-MIN106D | SQK-DCS109 + EXP-NBD104 | MinION | FAQ51879_pass_barcode03_  03abc106_254e4132_*****.fastq.gz |
| HeLa_directcDNA_replicate2_run1 | HeLa | Barcode4 | direct-cDNA | FLO-MIN106D | SQK-DCS109 + EXP-NBD104 | MinION | FAQ51879_pass_barcode04_  03abc106_254e4132_*****.fastq.gz |

The figures presented in the Supplementary Figure S1-S9 are associated to data used in Supplementary Table 1.

The * represents several fastq files generated per condition following sequencing. The fastq files can be accessed and downloaded in the following link:

<https://seafile.rlp.net/d/7a99b8b210e44eb9b70a/?p=%2FHEK_HeLa_Test_Upload%2FHEK_HeLa%2F20230323_1311_MN32609_FAQ51879_03abc106%2Ffastq_pass&mode=list>

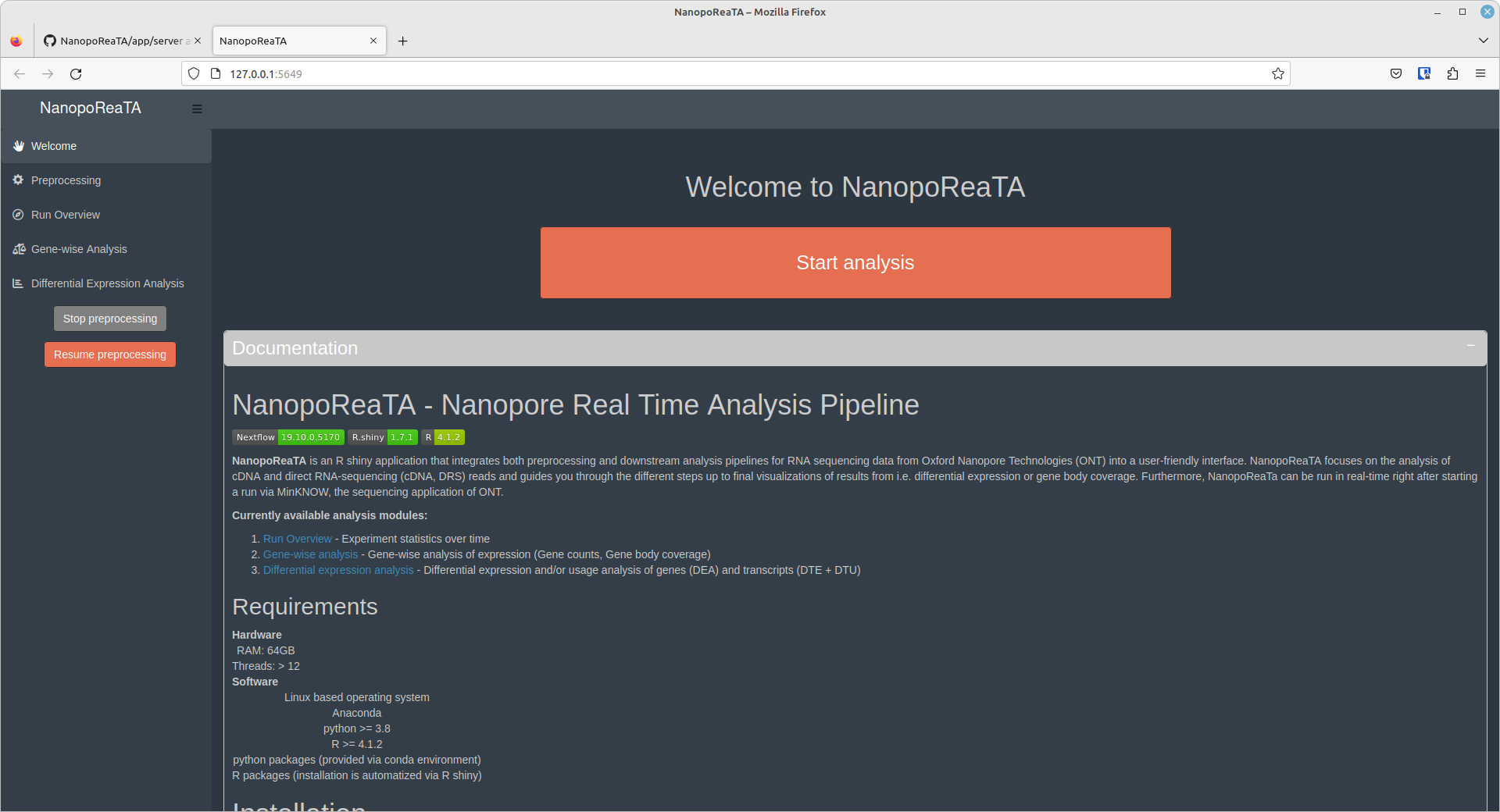

**Supplementary Figure S1 – Welcome tab.** The welcome page contains the github documentation of NanopoReaTA as well as the “**Start NanopoReaTA”** button to initiate the analysis pipeline.

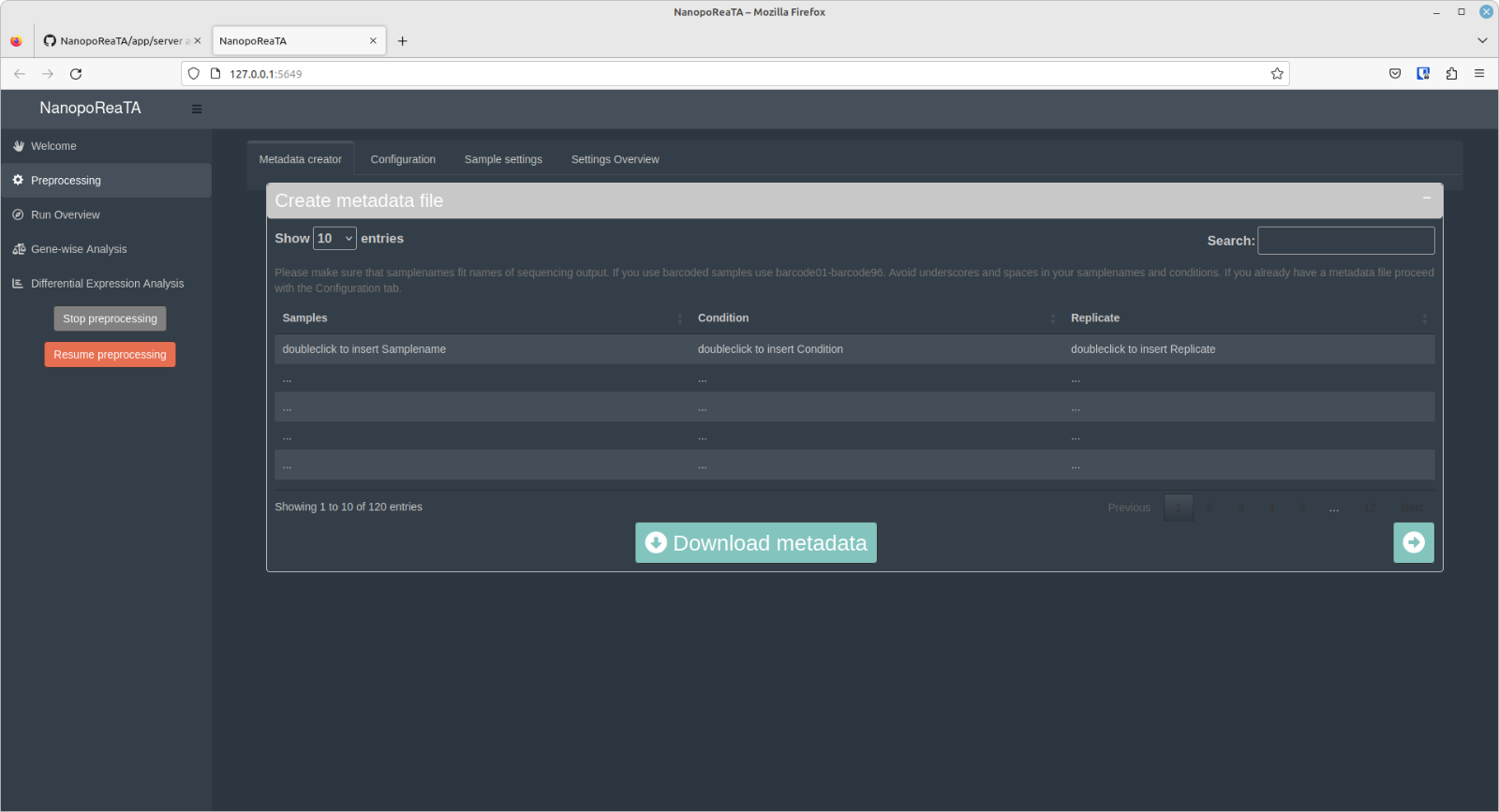

A

B

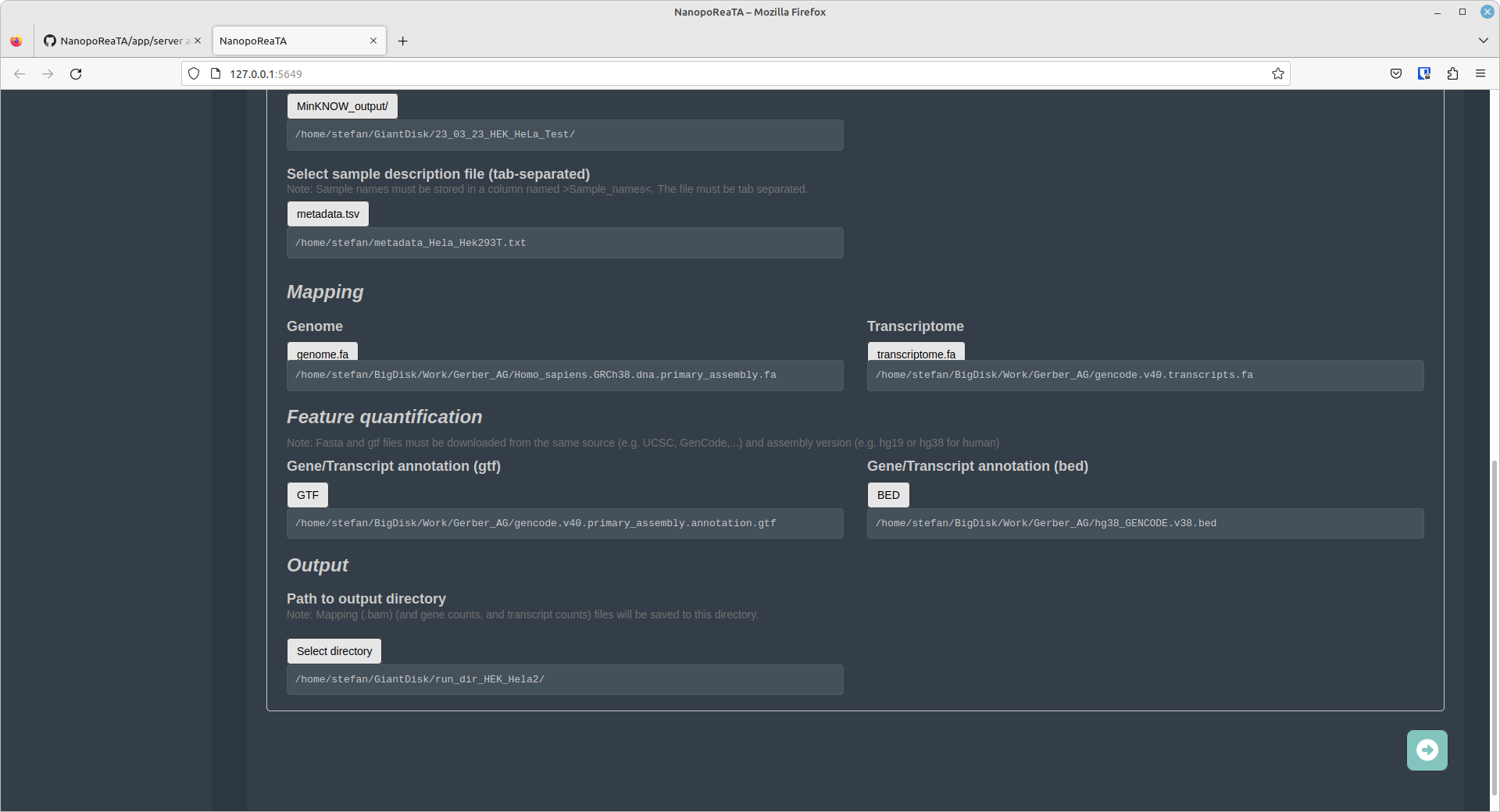

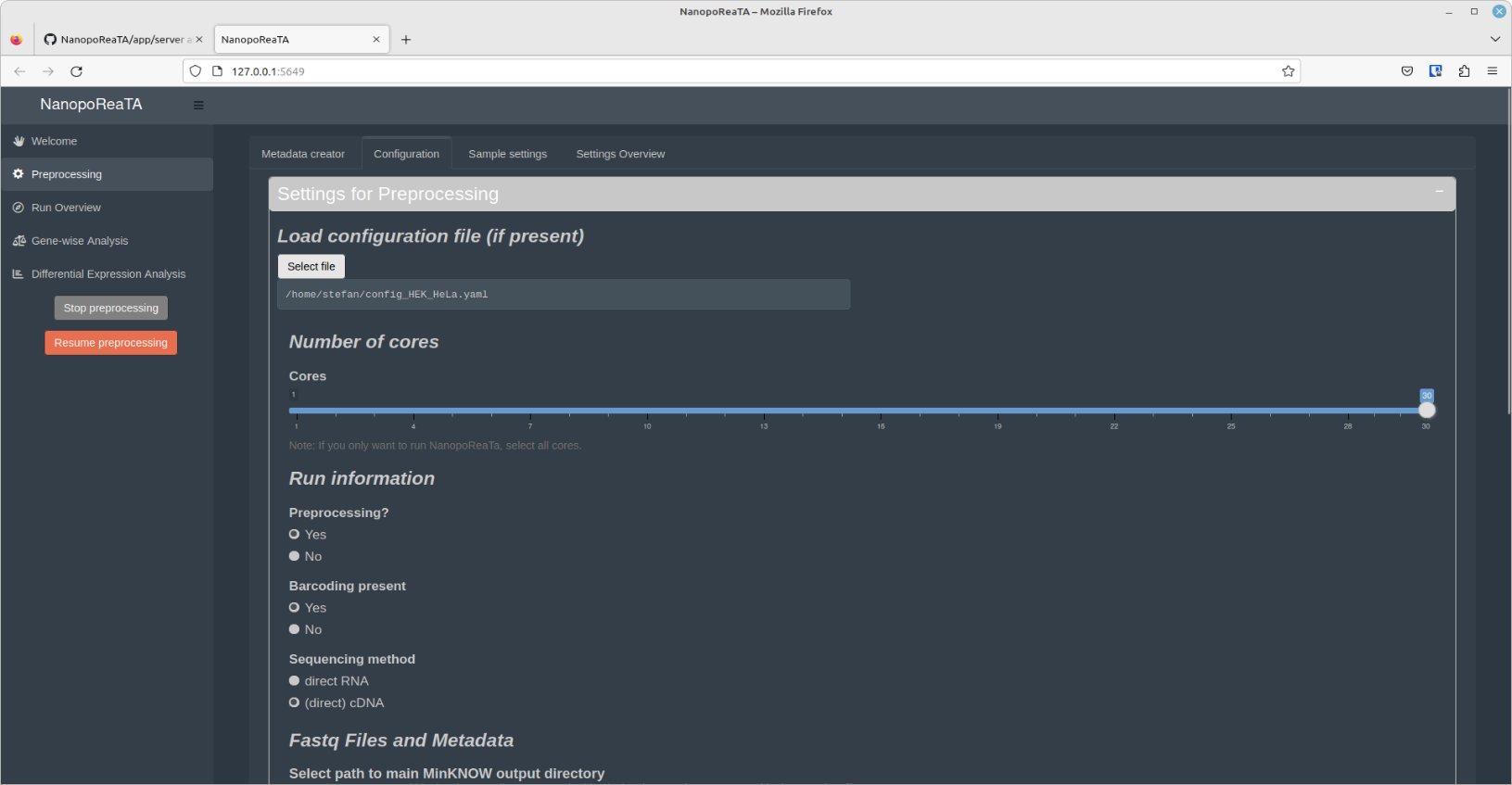

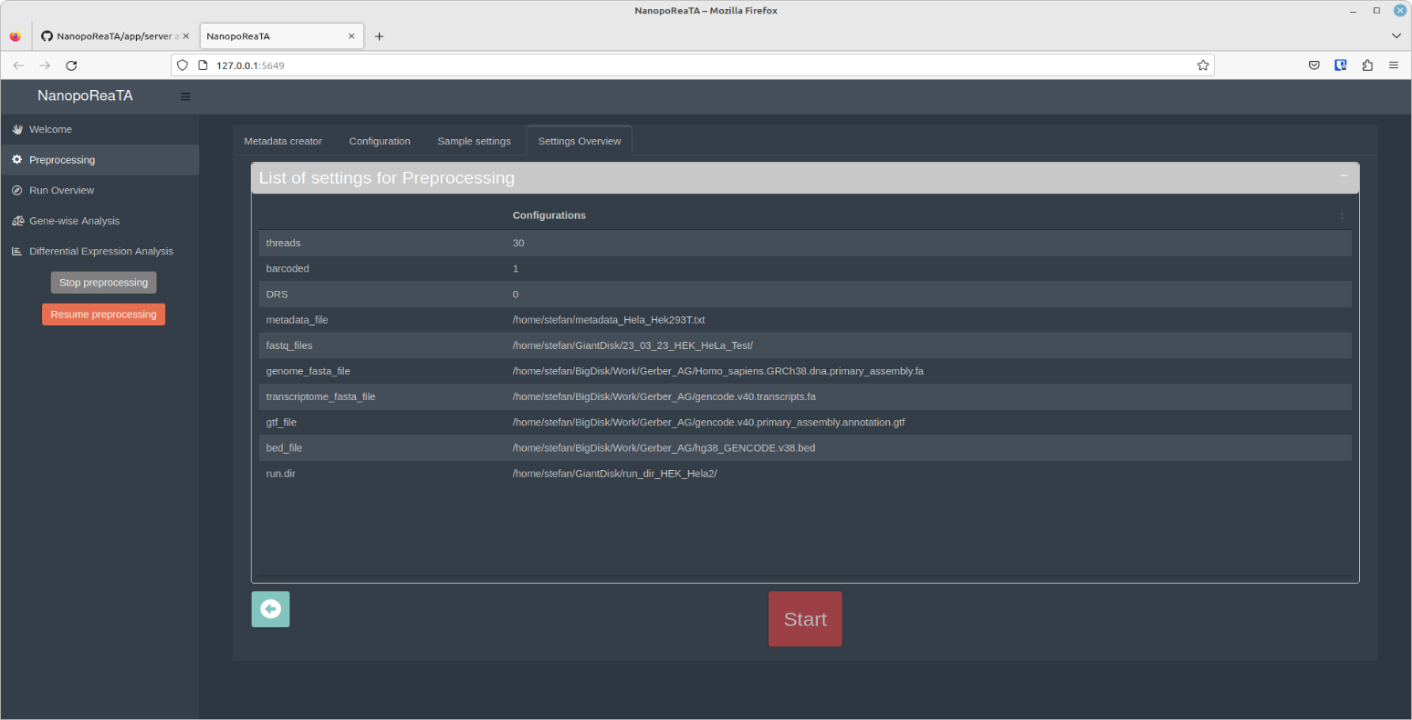

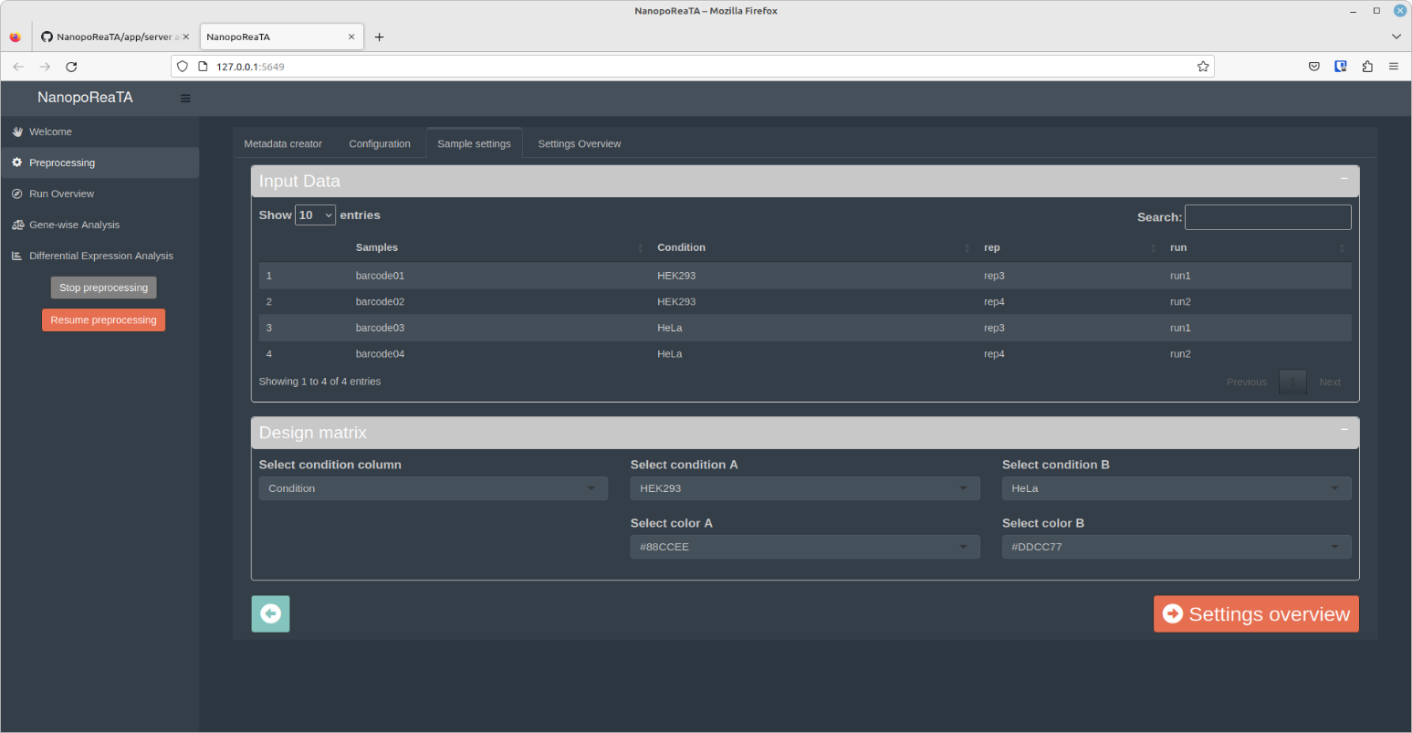

D

C

**Supplementary Figure S2 – Preprocessing tab.** The preprocessing tab provides a work frame that leads the user through input settings boxes to generate a complete configuration file that contains all information for preprocessing and downstream analyses. Instead of manual input, a configuration file containing all required information can be uploaded (an example configuration file can be found in the github repository). **A.** The “Metadata Creator” allows to create a metadata table that includes all necessary information about the samples for the pipeline configuration. **B.** The “Configuration" tab contains the setup for all user inputs needed for following analyses steps. Here the user can select the number of cores to be used, if preprocessing should be applied and if samples are barcoded. In case of config file upload, the information will be automatically put into respective boxes shown at this page. Once the data is checked, the white button on the bottom right will confirm the use of the config file (check config.yaml for correct parameters settings) **C.** The “Sample settings” table shows the input metadata table where the user can check all information is loaded correctly. For pairwise comparison, the user needs to select one of the columns containing two conditions that will be compared in further analyses. For visualizations, the user can change the color-coding for each condition here. **D.** The “**Settings overview” tab** contains the final configuration overview where the input settings can be finally checked. If the parameters are correct, the user can start the preprocessing pipeline by clicking the **Start preprocessing** button. Otherwise, the user can rearrange the settings by going back to the configuration tab.

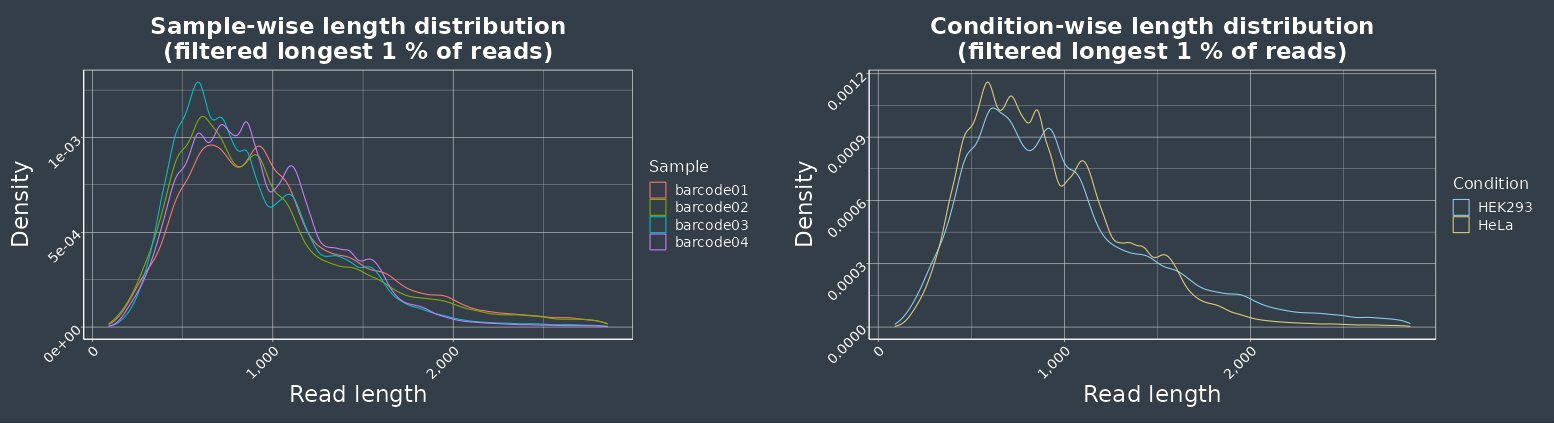

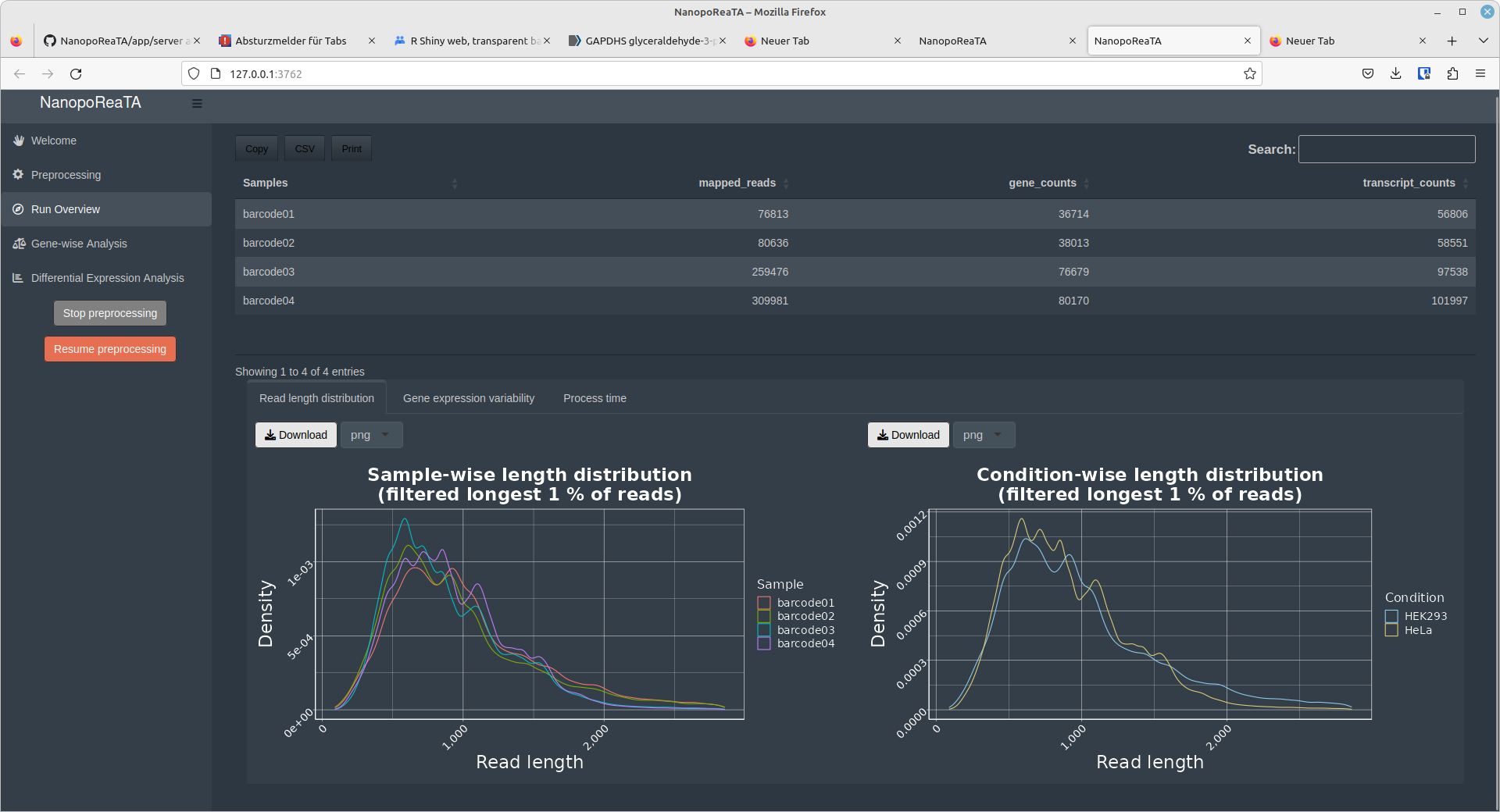

C

B

A

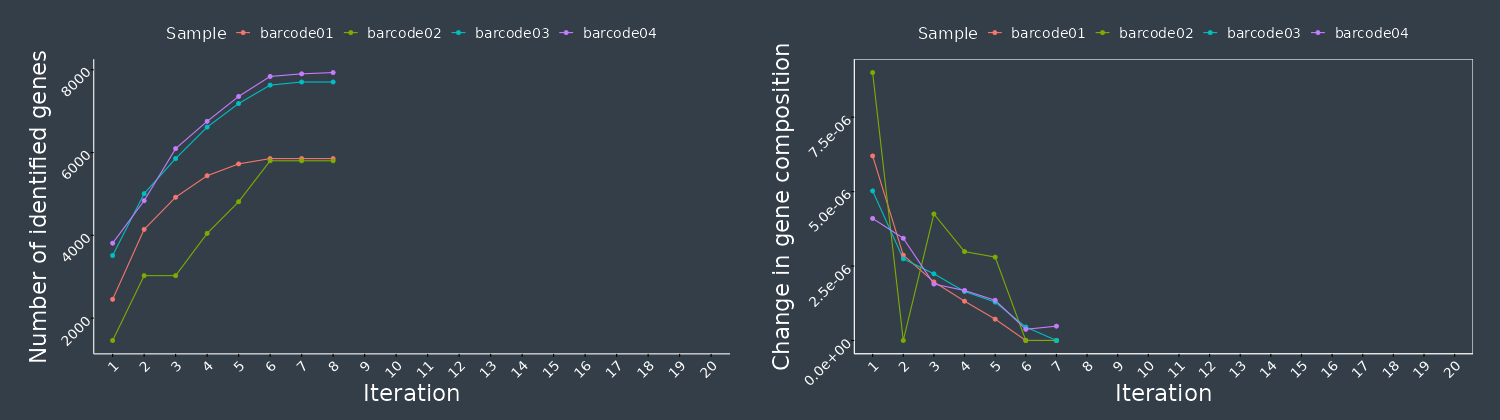

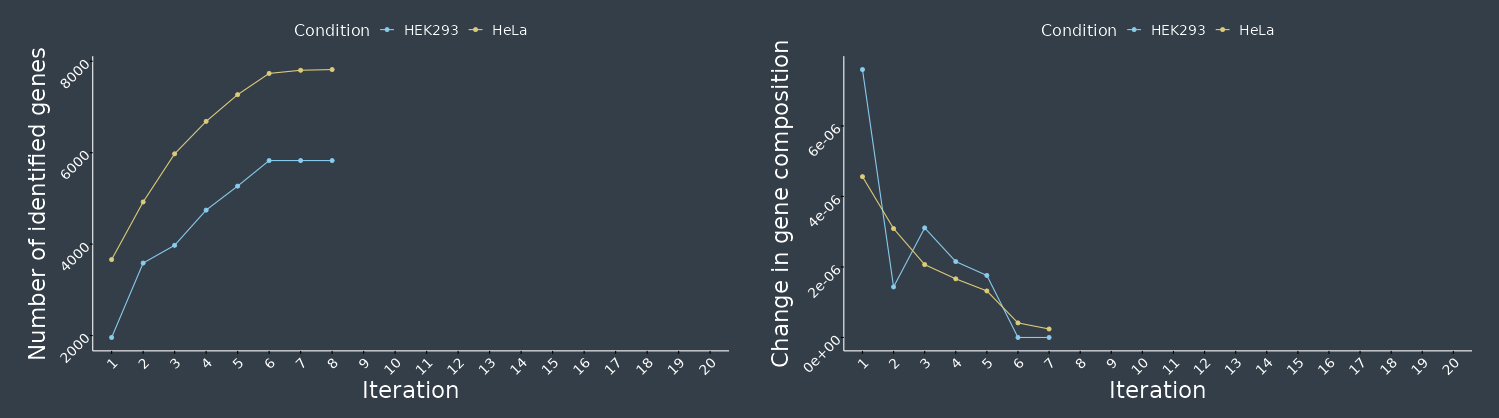

Figure continues next page.

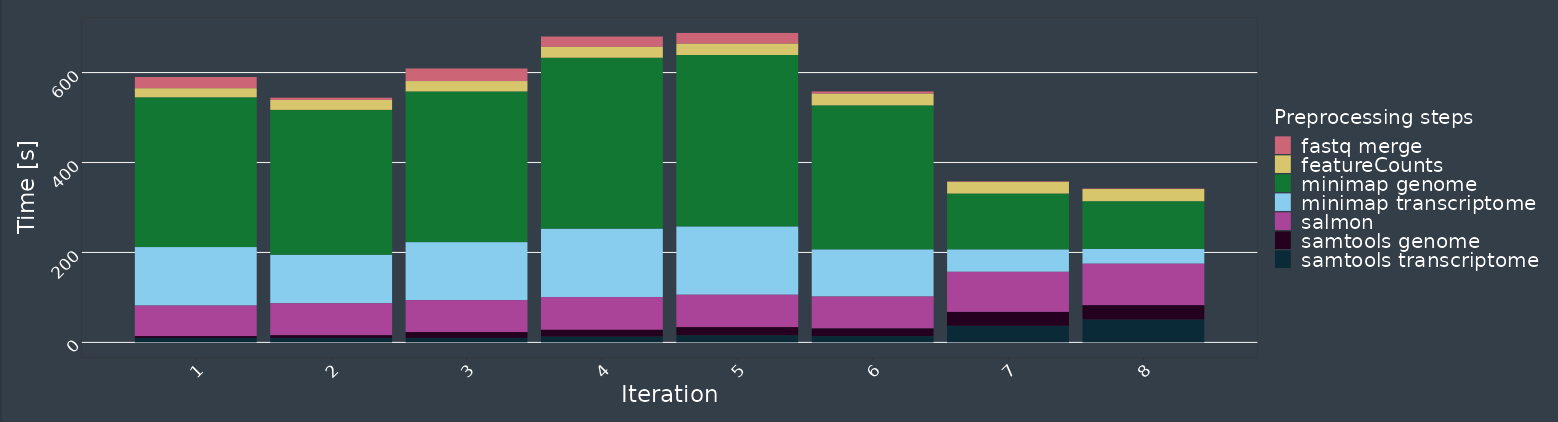

D

**Supplementary Figure S3. Run Overview.** **A.** The data table at the top shows the number of mapped genes (minimap2), gene counts (featureCounts) and transcriptome (salmon) counts. The counts are provided for each sample respectively. **B.** Read length distribution. Read length distributions are presented for respective samples or selected conditions. The read length information is extracted directly from the fastq files (passed_reads). **C.** Gene count variability. The plot on the left presents the number of genes detected per iteration for individual samples (top) or selected conditions (bottom). The information is extracted from the output count table of feature counts. The plot on the right presents the deviation of relative gene abundancy compared to the last iteration. Such measure reports the change of gene abundancy variability within a single sample revealing the stabilization of relative gene abundancies throughout the ongoing sequencing. **D.** Process time. These plots represent the run time of each tool during the preprocessing per iteration. (See “Run Overview” in <https://github.com/AnWiercze/NanopoReaTA>).

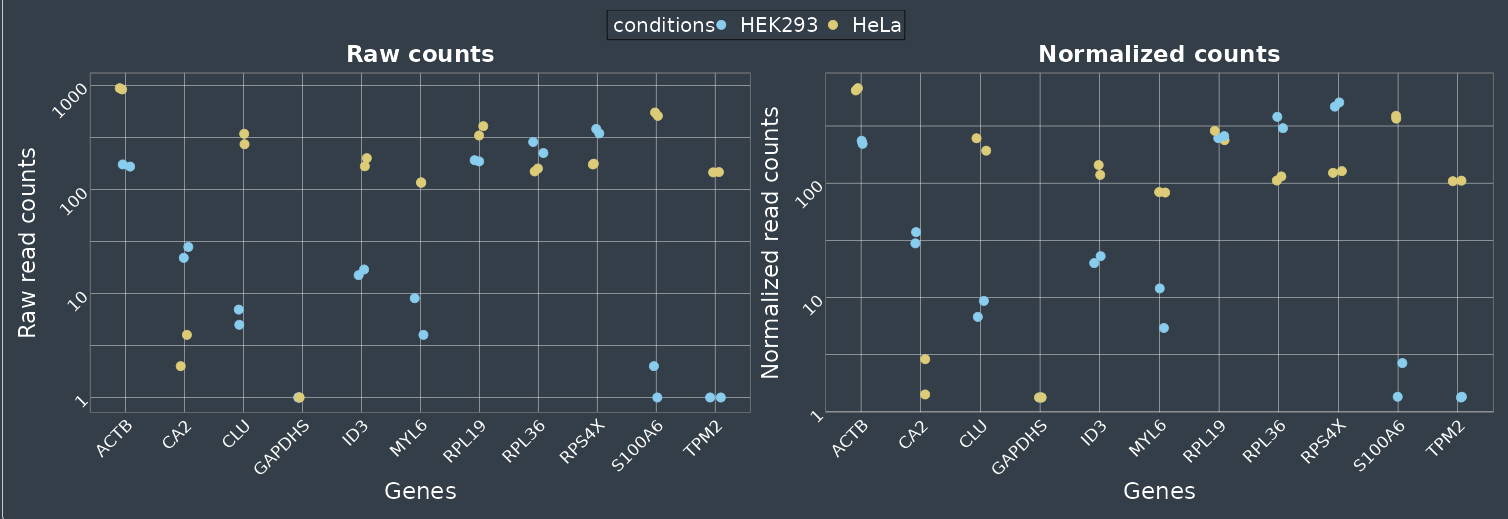

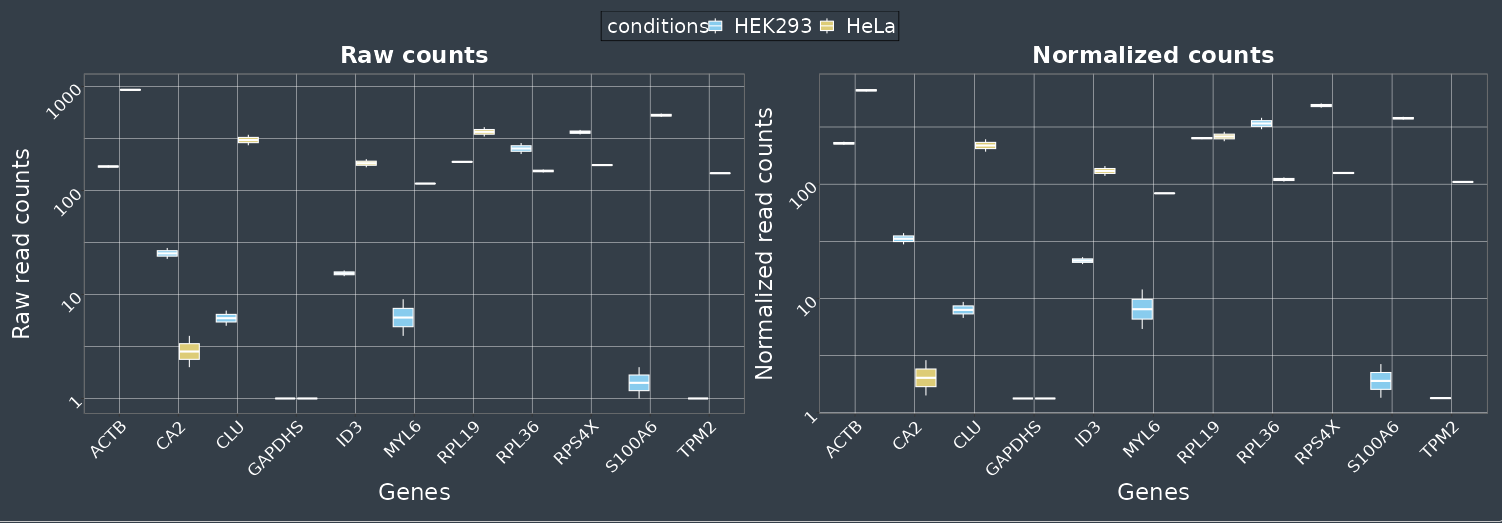

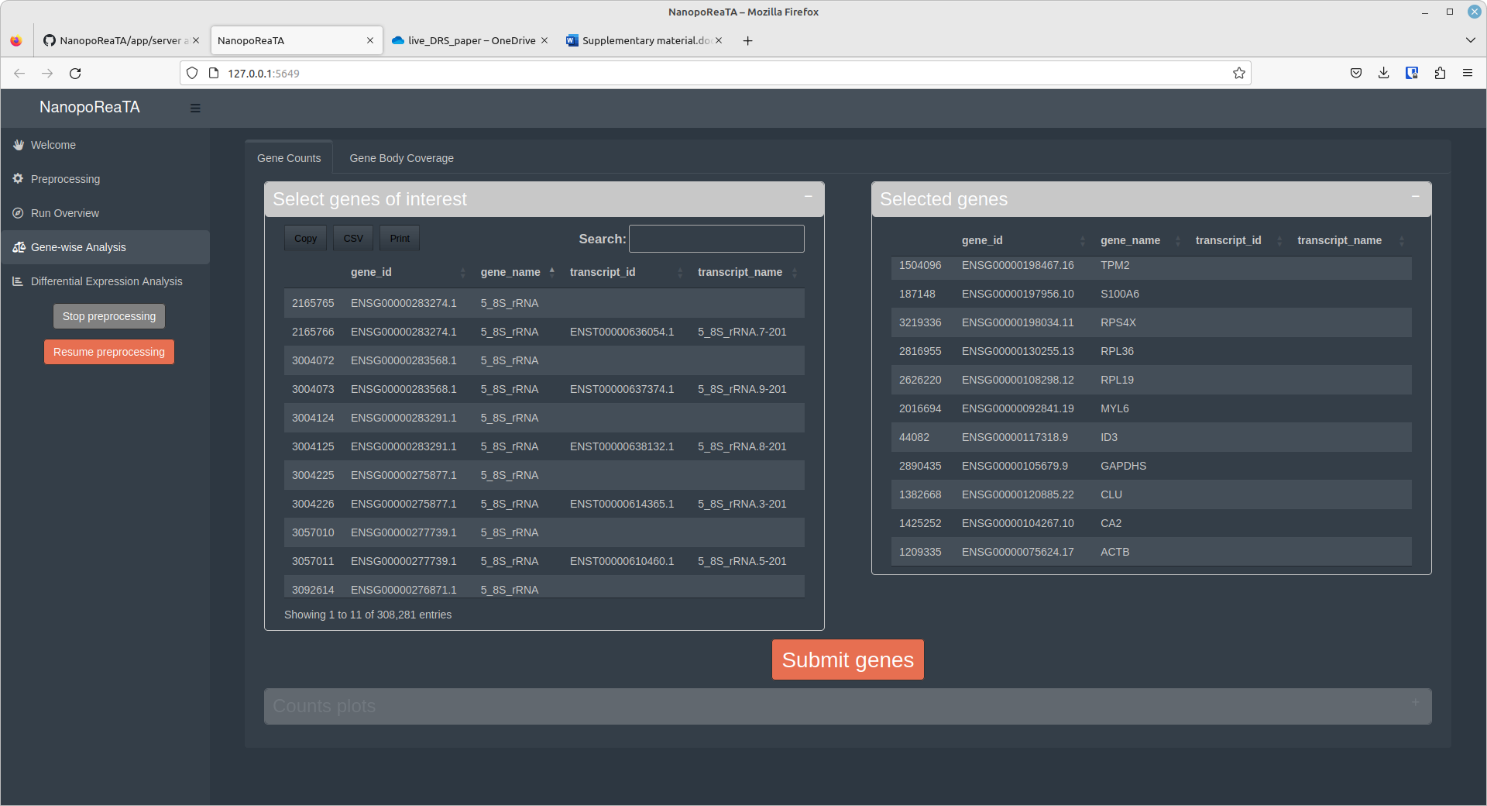

Bv

Av

Figure continues next page.

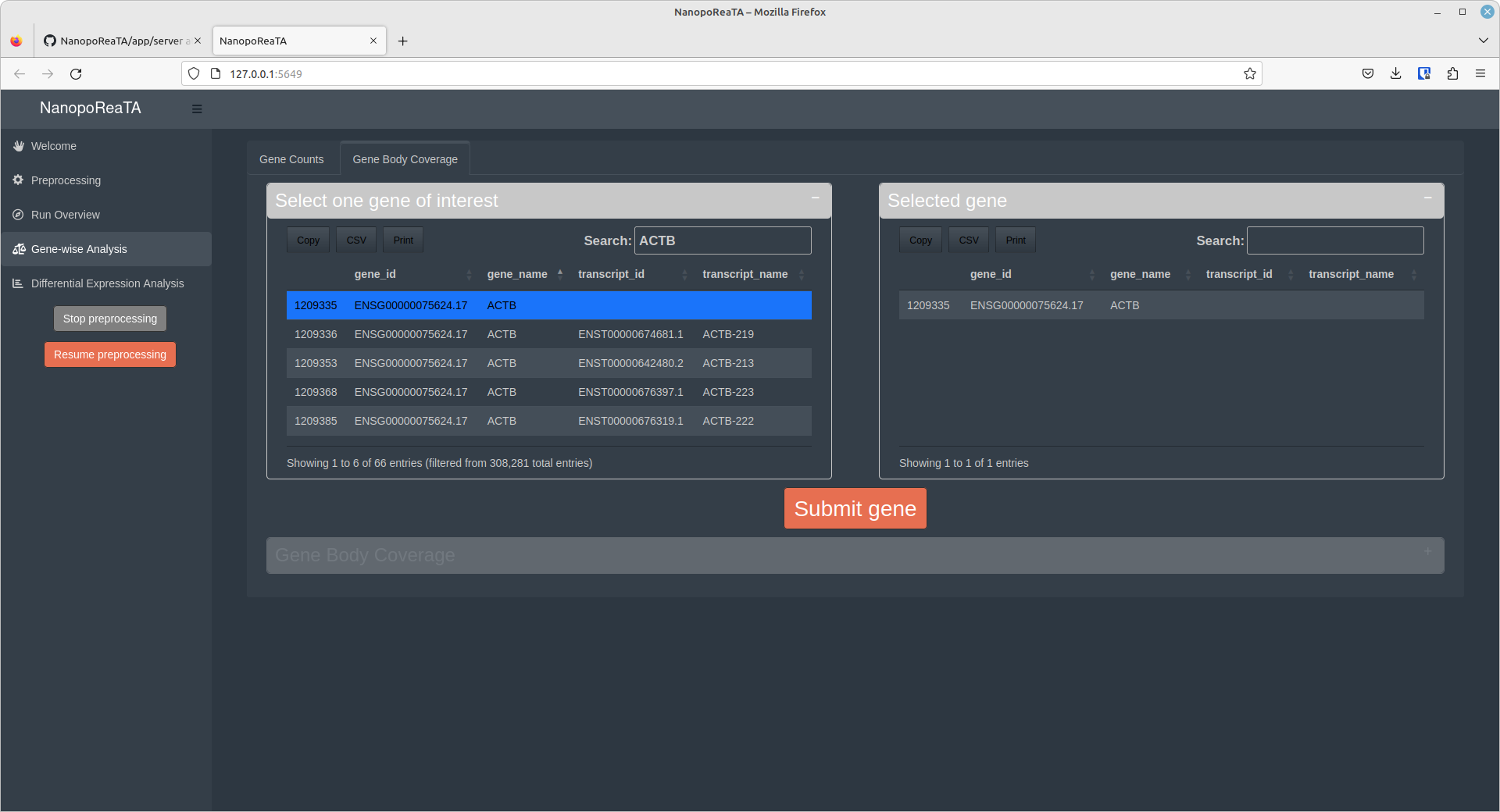

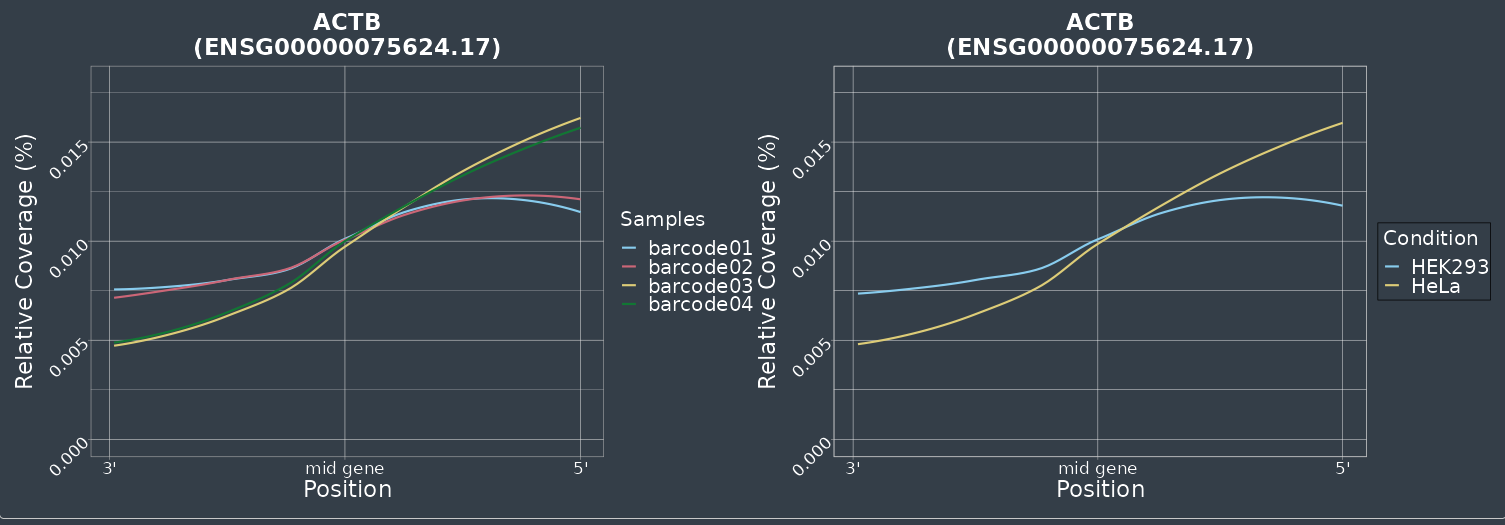

D

C

**Supplementary Figure S4- Gene-wise analysis tab.** This tab presents the expression levels and the gene body coverage of specific genes of interest. Be aware that at least two samples per condition have to be considered in order to use this functionality. **A.** The table on the left presents all the genes annotated in the loaded GTF file. Individual or several genes of interest can be selected via double click on the table entry. Selected genes will appear in the table at the right and by clicking the “submit genes” button, the analysis will start. **B.** The median of ratio normalization of the selected genes will be performed via DESeq2 and plot the expression of the selected genes as Dot- or Boxplot. The plots present the raw counts (left) and normalized counts (right) (See “Gene-wise analysis - Gene counts” in <https://github.com/AnWiercze/NanopoReaTA>). **C.** Gene body coverage allows the analysis of an individual gene selected from the table on the left. The selected gene will appear in the table at the right and by clicking the “submit gene” button, the analysis will start. The plots represent the percentage of coverage for exon-percentiles for single samples (left) and for selected conditions (right).

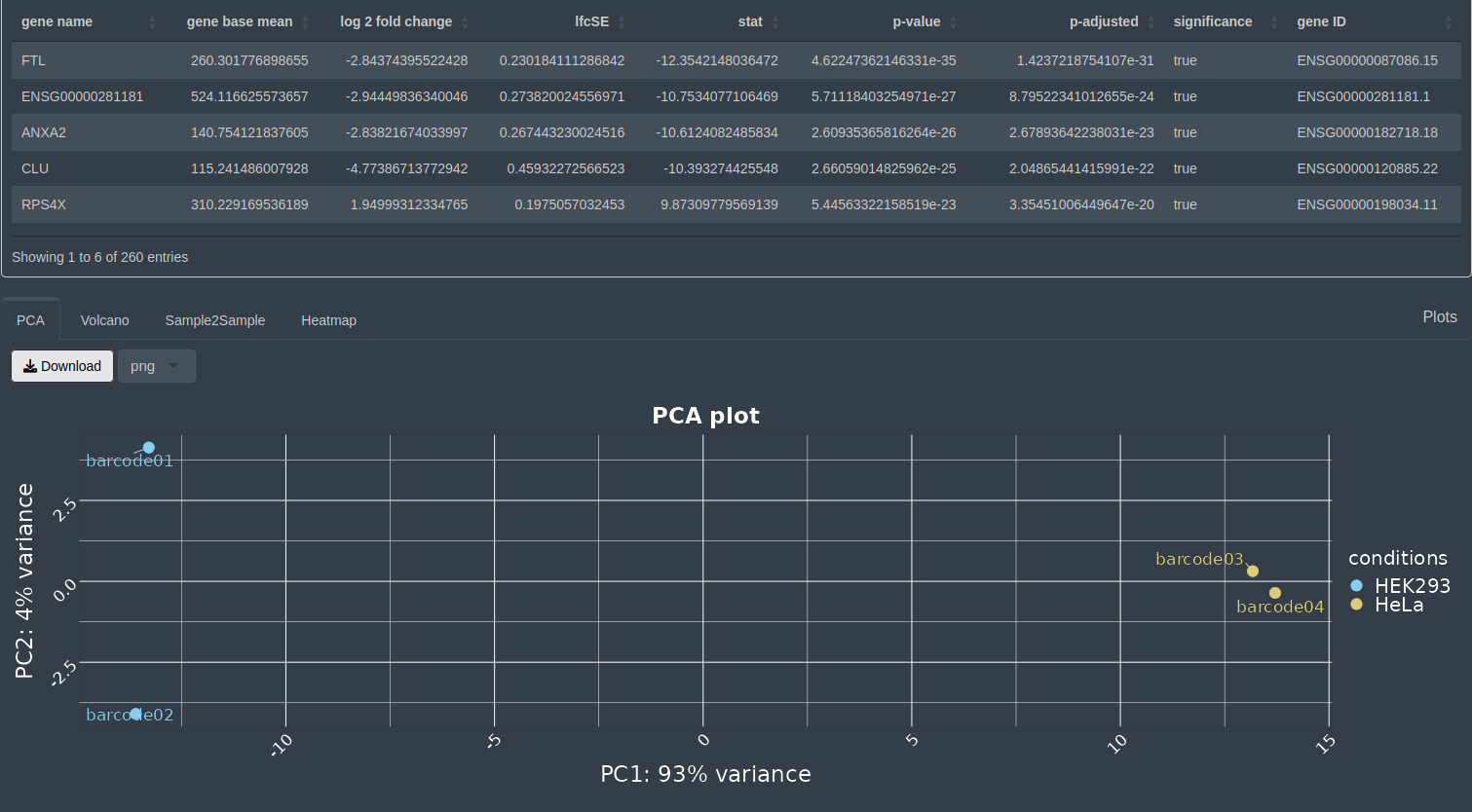

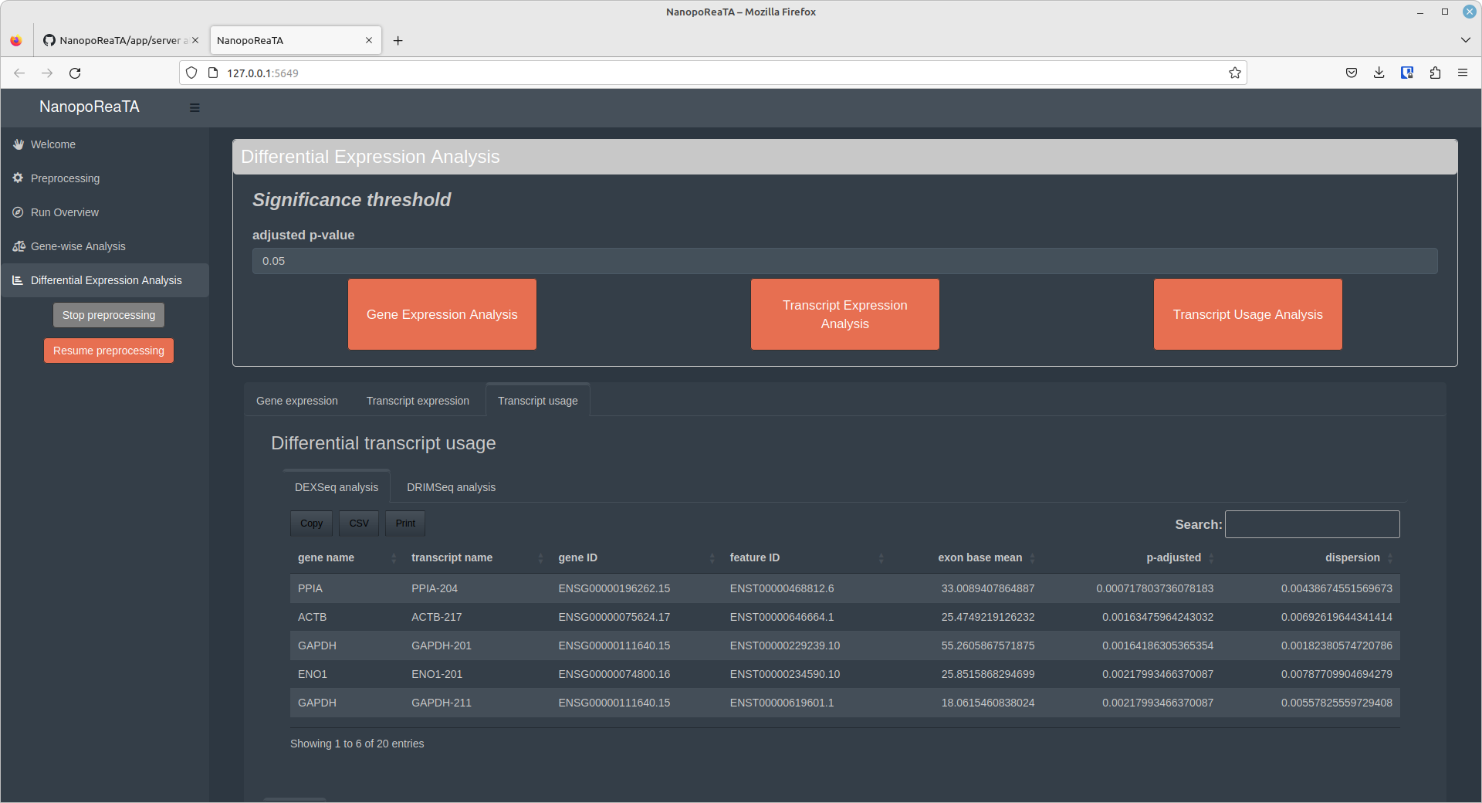

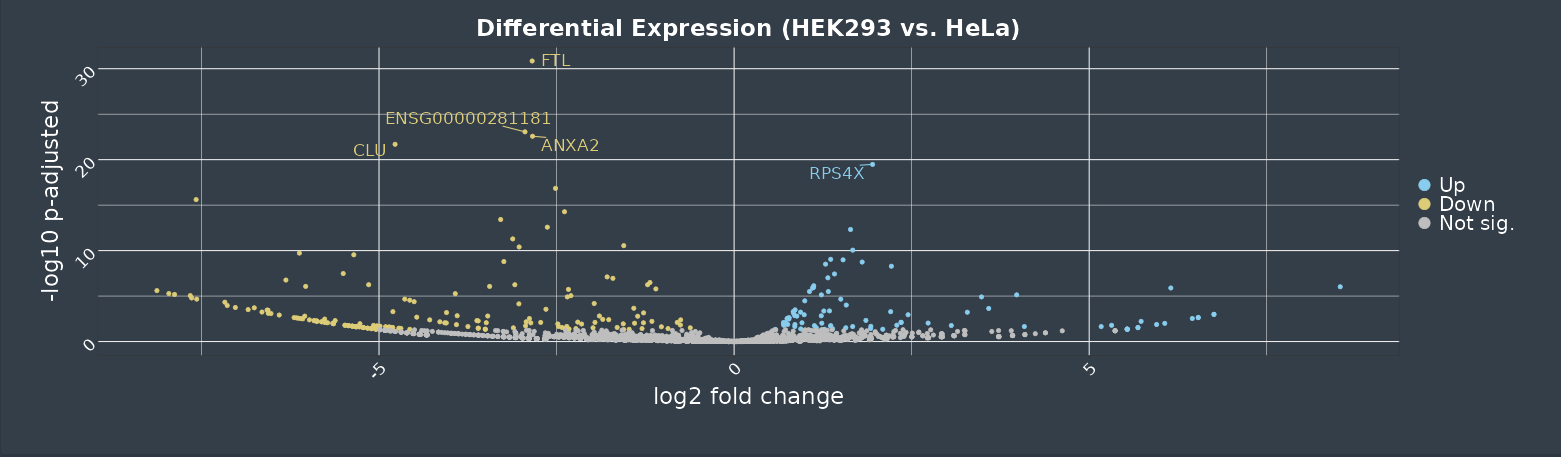

C

B

A

Figure continues next page.

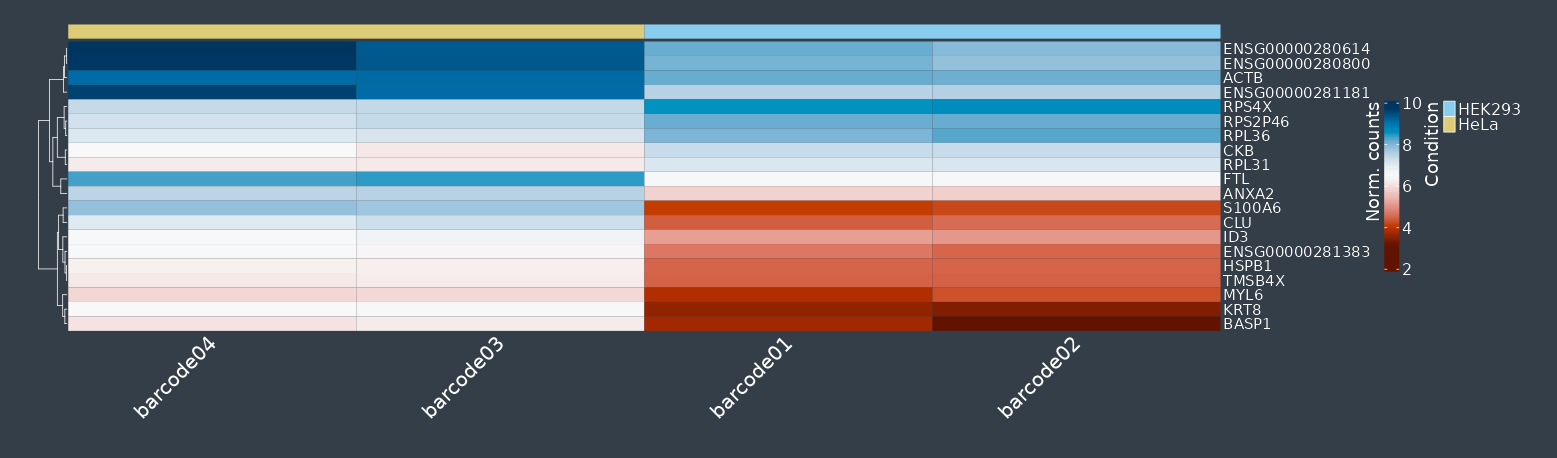

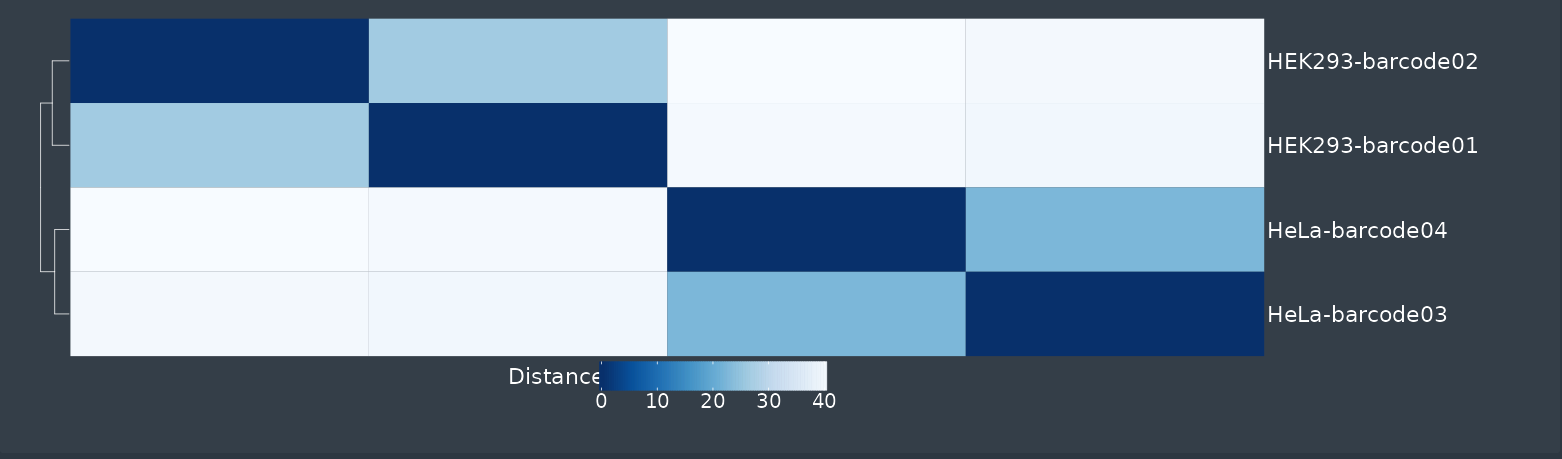

E

D

**Supplementary Figure S5- Differential expression analysis – gene and transcript level.** For expression analysis, following each iteration, the user can press on the “Gene expression analysis” or “Transcript expression analysis” button which updates the DESeq2 analysis step for the iteration. Note: Sequencing should run long enough to show low inner variabilities in the expression pattern. After pressing the button a differential expression analysis is executed. This may take around 5 minutes, depending on the size of the dataset. Following the analysis, several datasets and plots can be retrieved including: **A.** Table of all differential expressed genes **B.** PCA plot **C.** Volcano Plot (DGE)) **D.** Sample-2-sample plot **E.** Heatmap of the top 20 differential expressed genes (based on p-adjusted). The tables and plots can be downloaded as csv or png/pdf, respectively. The figure shows results of differential gene expression, which are analogue to transcript expression analysis. (See “Differential Expression Analysis” in <https://github.com/AnWiercze/NanopoReaTA>)

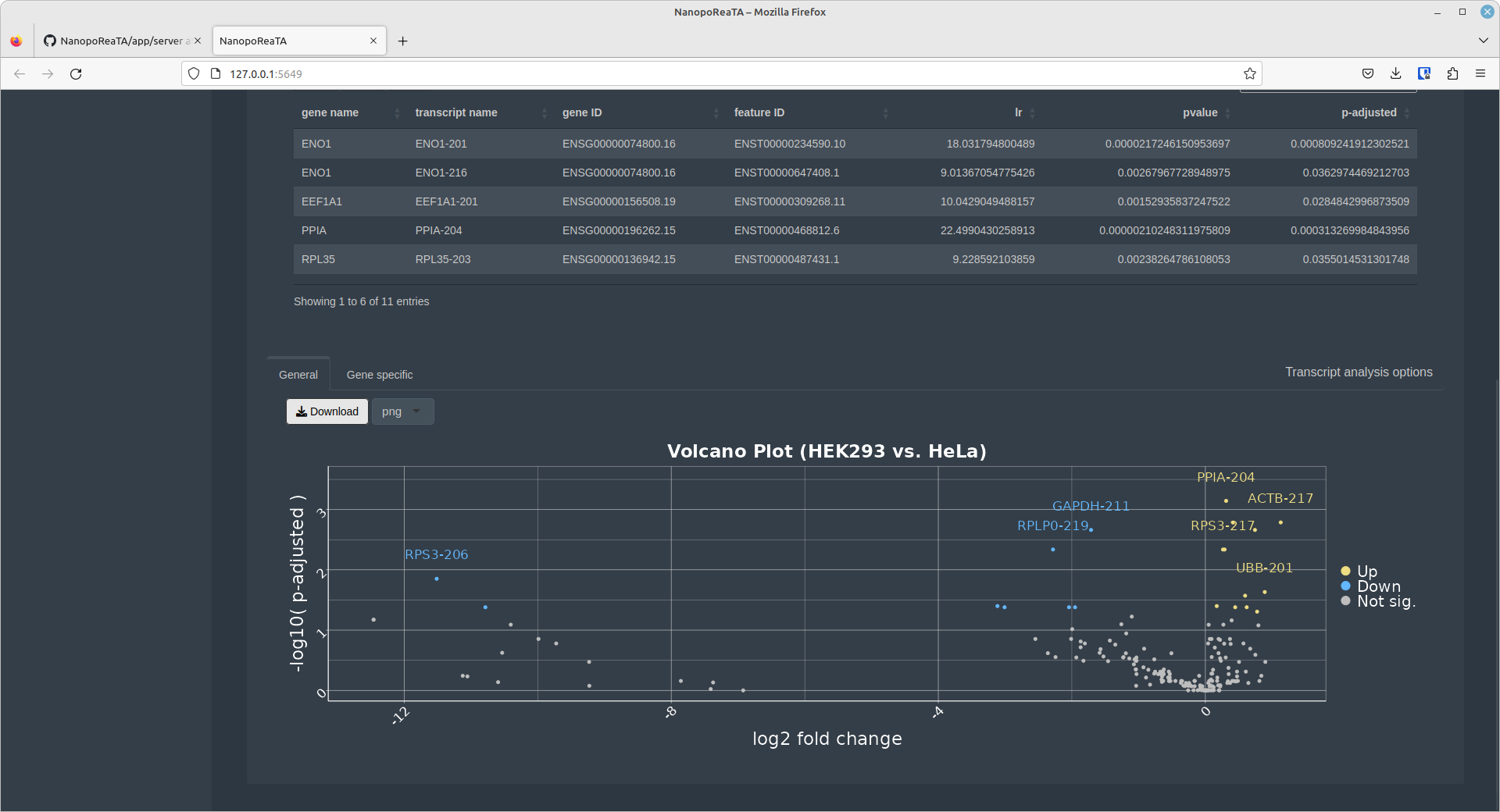

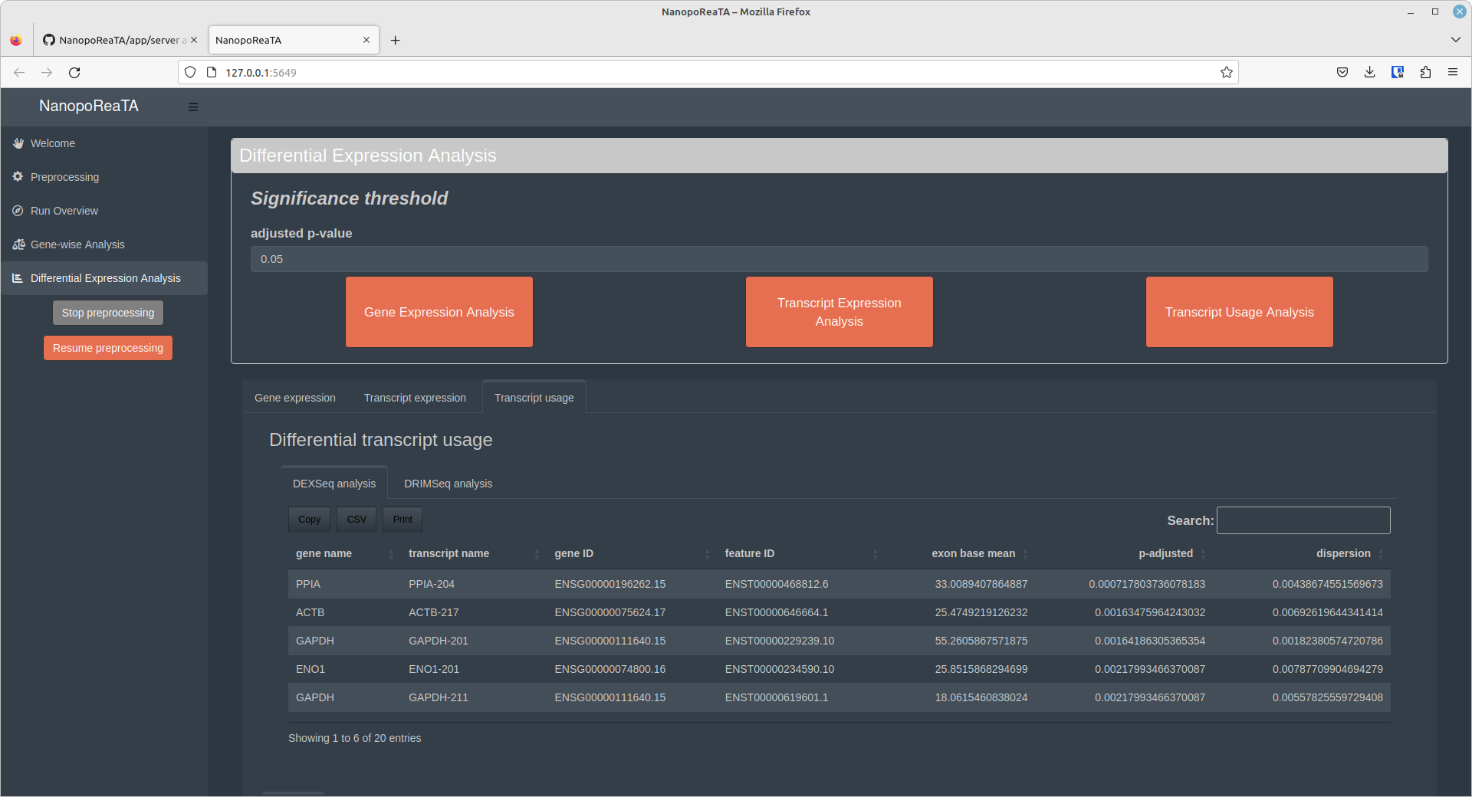

B

A

Figure continues next page.

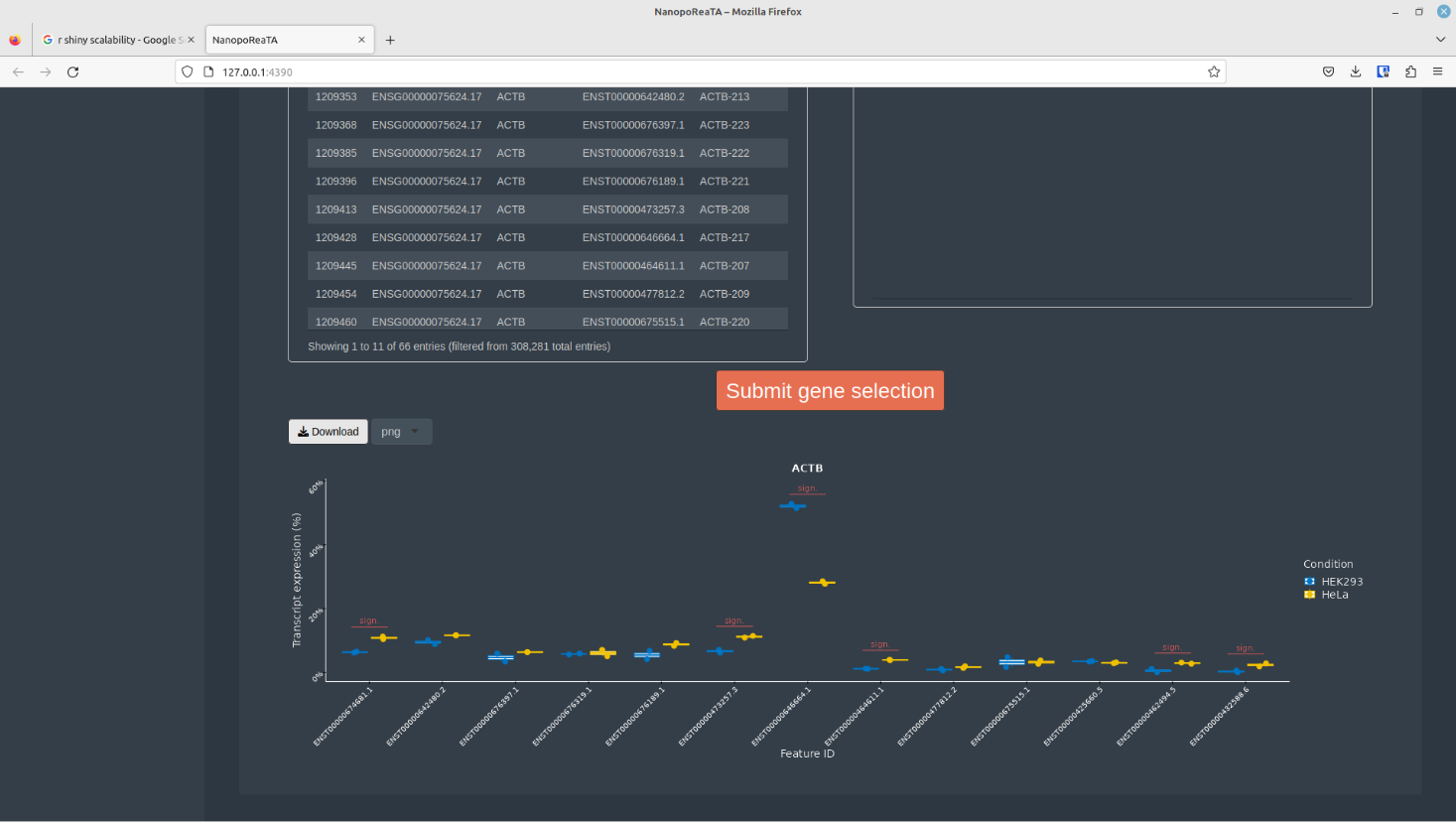

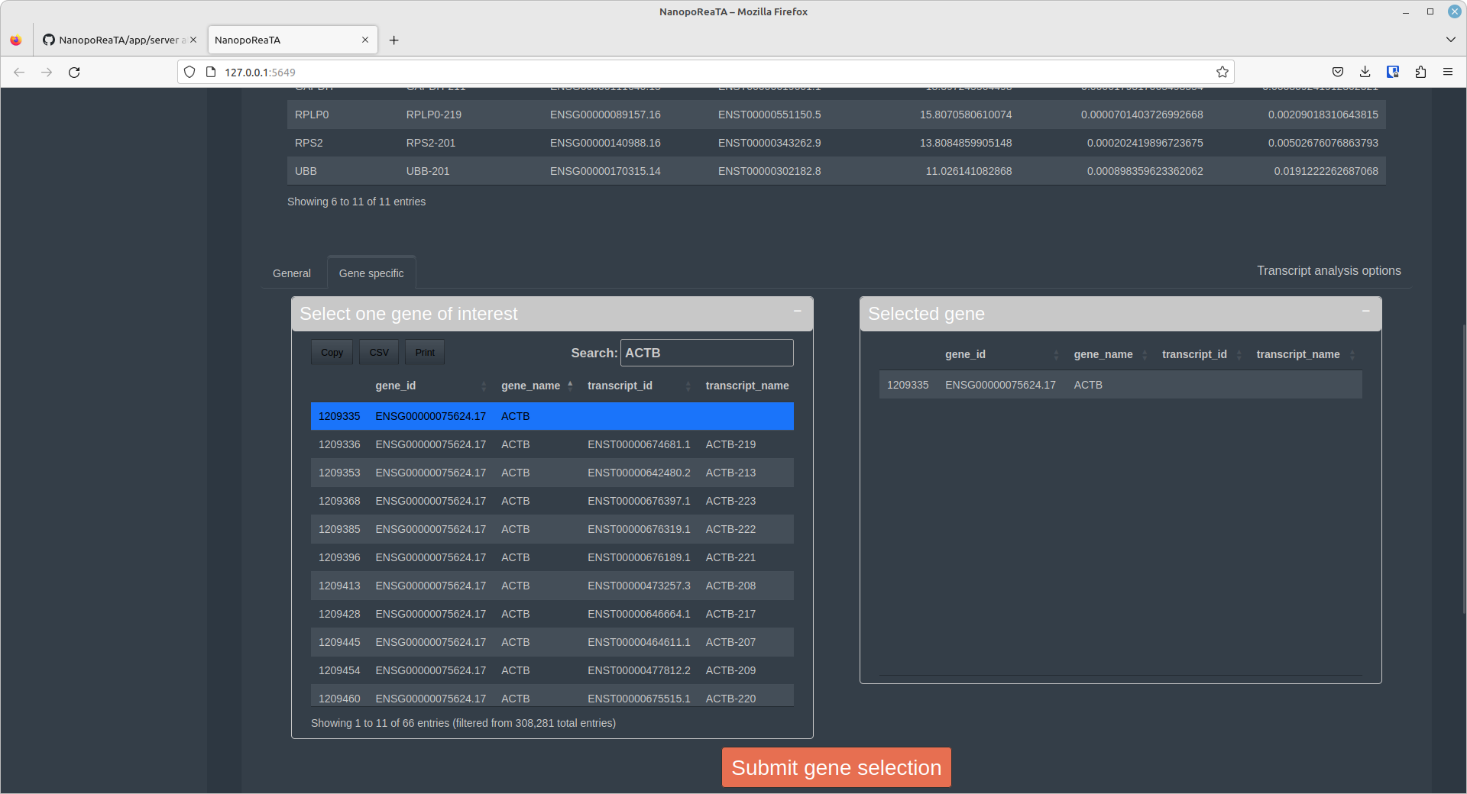

C

**Supplementary Figure S6- Differential transcript usage analysis.** For differential transcript usage analysis, following each iteration, the user can press on the “Transcript usage analysis” button which updates the count-files for the transcripts using DRIM-Seq and DEXSeq pipelines. Note: Sequencing should run long enough to show low inner variabilities in the expression pattern. After pressing the button, a differential expression analysis is executed. This may take around 5 minutes, depending on the size of the dataset. Following the analysis, several datasets and plots can be retrieved including **A.** Table of all differentially used transcripts (DRIMSeq and DEXSeq analysis tables) **B.** Under differential transcript usage “General” tab, a volcano plot shows the most significant differentially used transcripts **C.** Under the differential transcript usage “Gene specific” tab, an individual gene can be selected for distinct annotated transcript variant analysis. (See “Differential Expression Analysis - Gene-level analysis (DGE with DESeq2” in <https://github.com/AnWiercze/NanopoReaTA>)

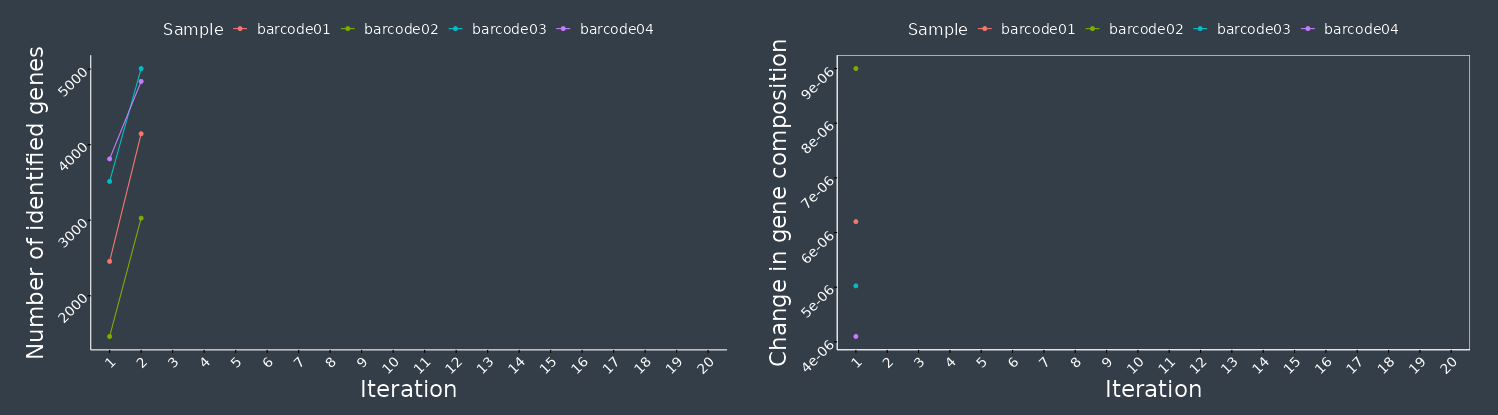

A

Iteration 2

B

Iteration 8

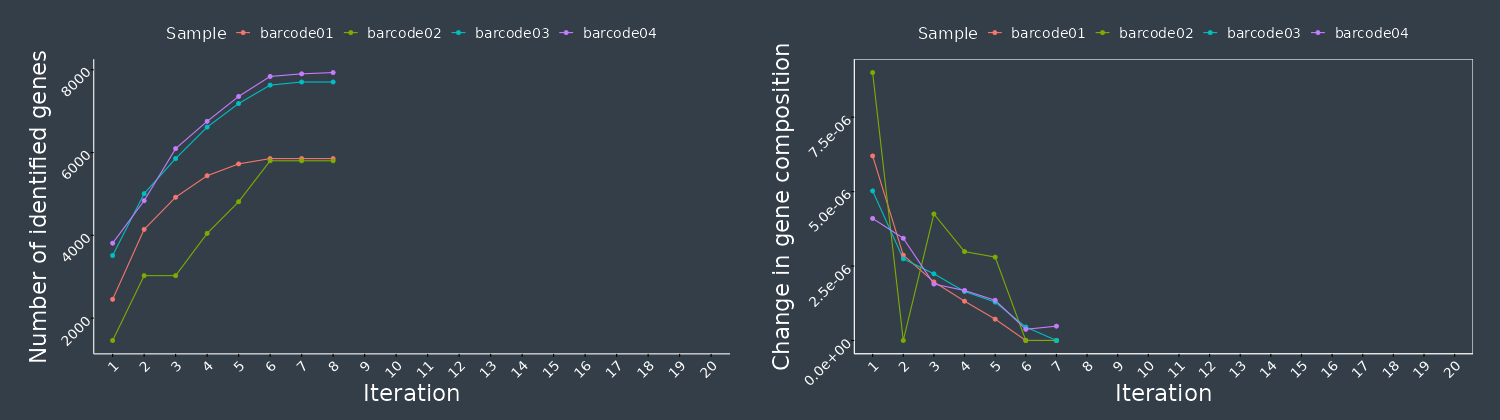

Figure continues next page.

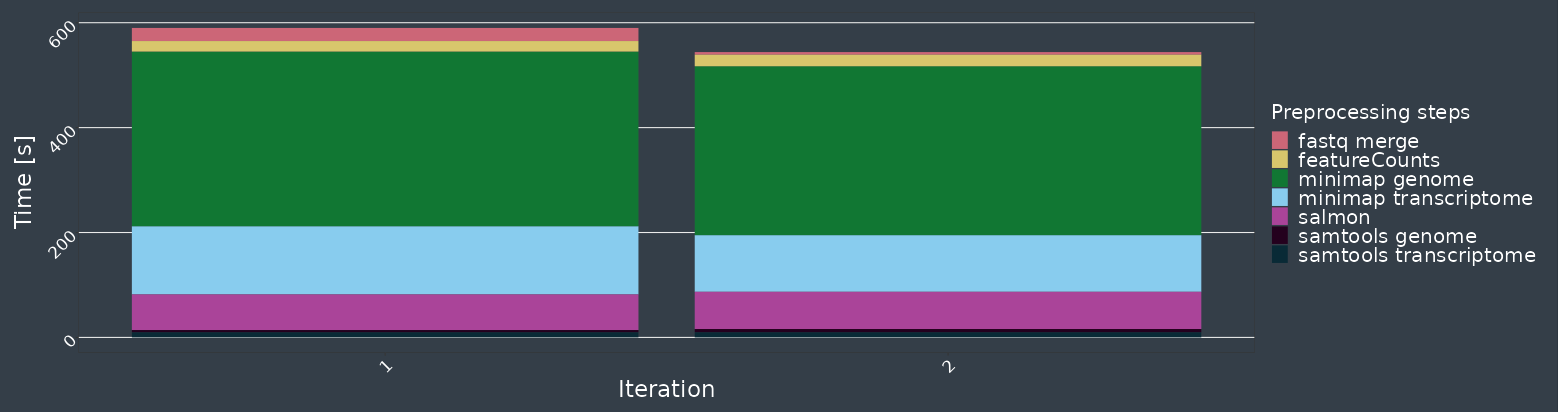

C

Iteration 2

D

Iteration 8

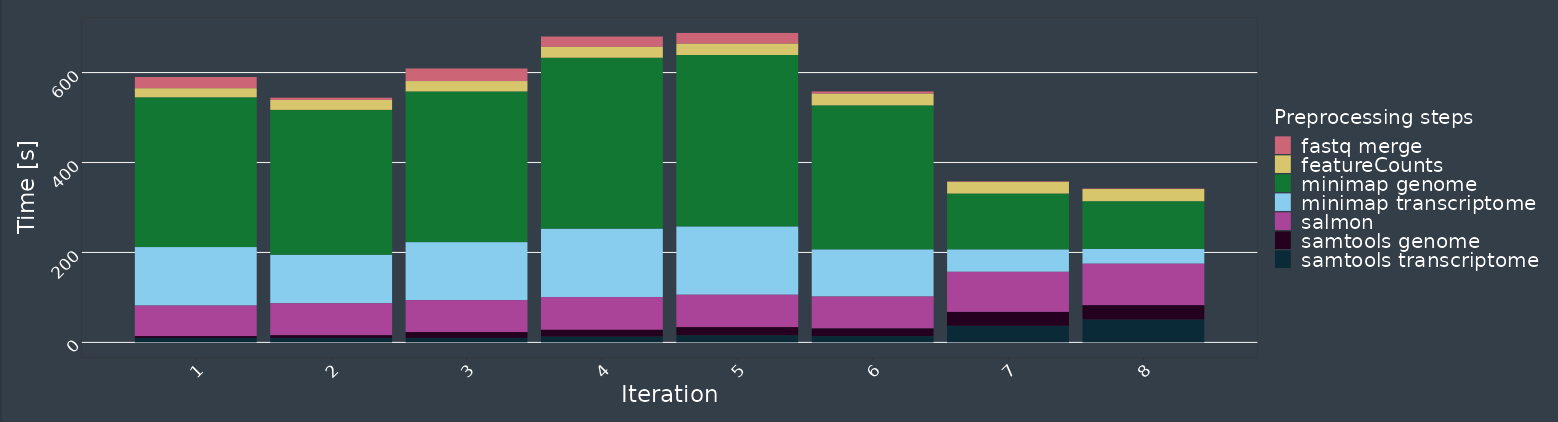

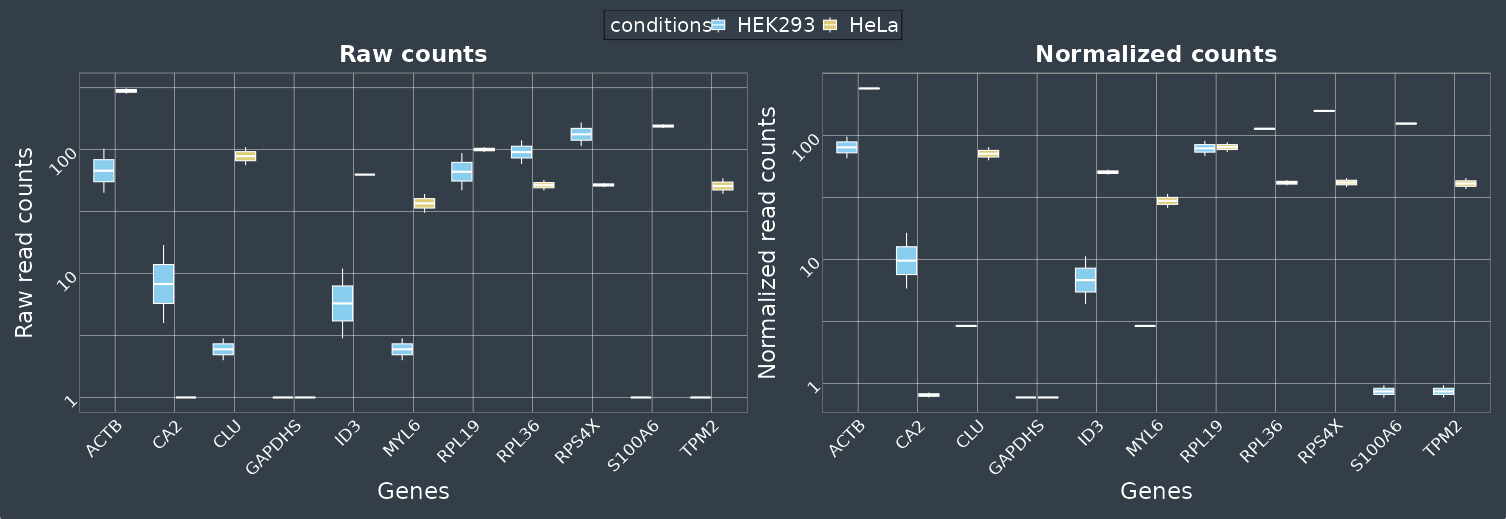

E

Iteration 2

F

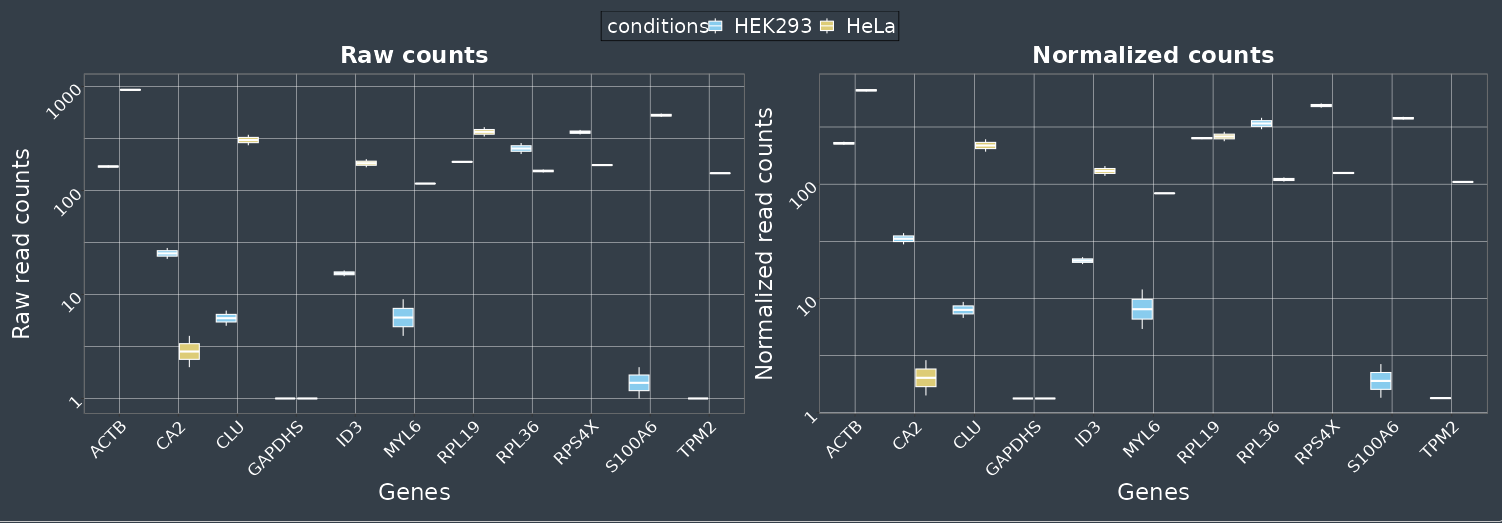

Iteration 8

**Supplementary Figure S7- Comparison of “Run overview” over time.** Iteration 2 (**A,C,E**) represents early sequencing state whereas iteration 8 (**B,D,F**) represents advanced sequencing state. **A-B**. Changes in “gene expression variability” demonstrate the increase detection of gene count overtime (left) and stabilization of gene abundancy variability (right) through time. **C-D.** Changes in “Process time” reveal the usage of each tool during the preprocessing through each iteration. **E-F.** Changes in “individual gene count” represent the expression of the selected genes (raw counts on the left and normalized counts on the right) throughout the sequencing run.

A

Iteration 8

Iteration 2

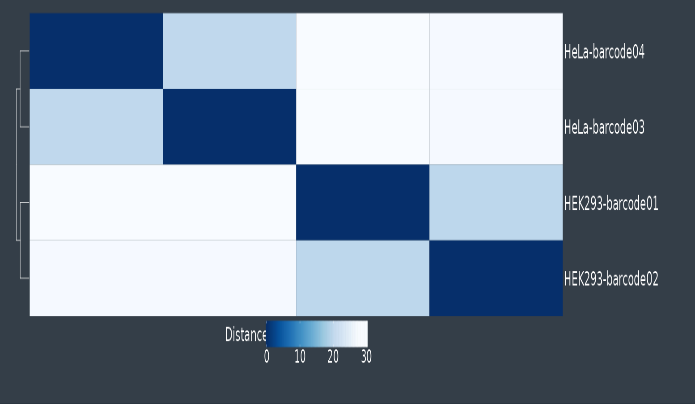

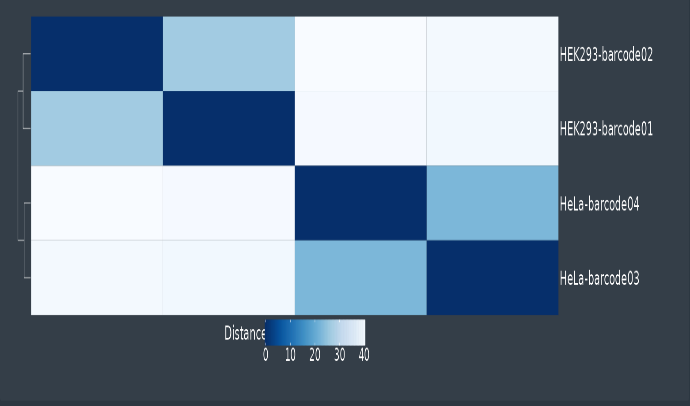

B

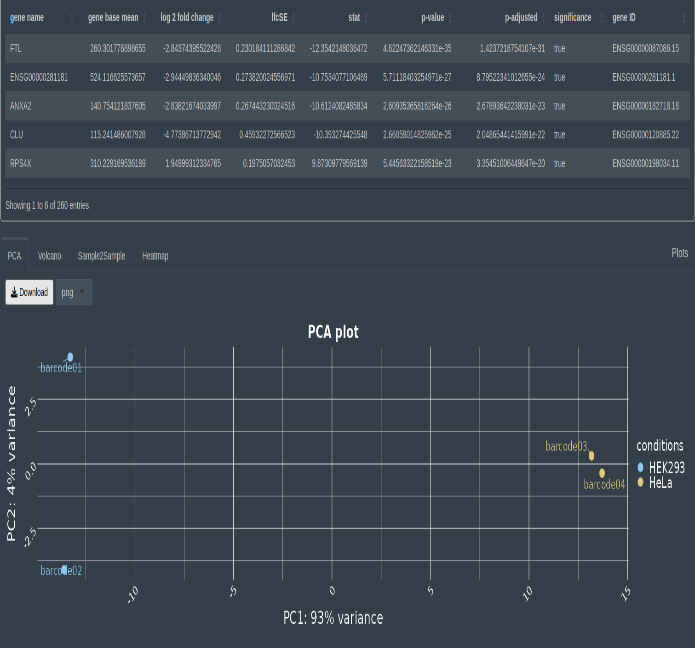

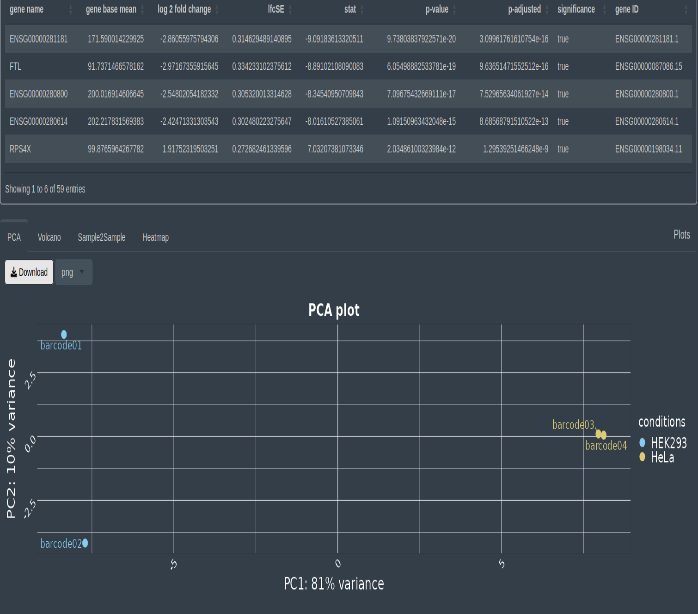

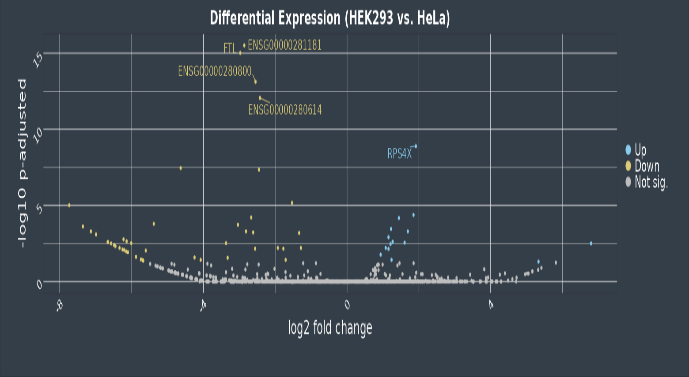

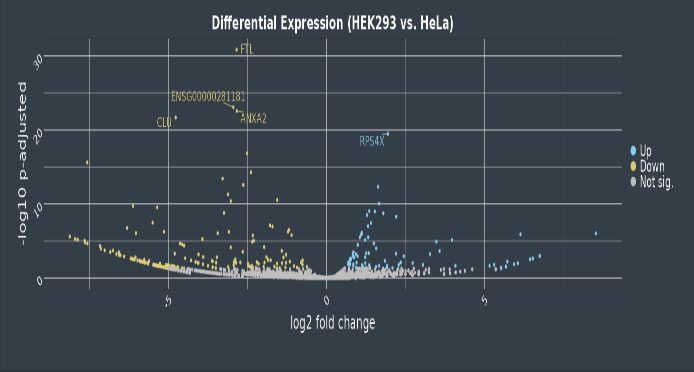

C

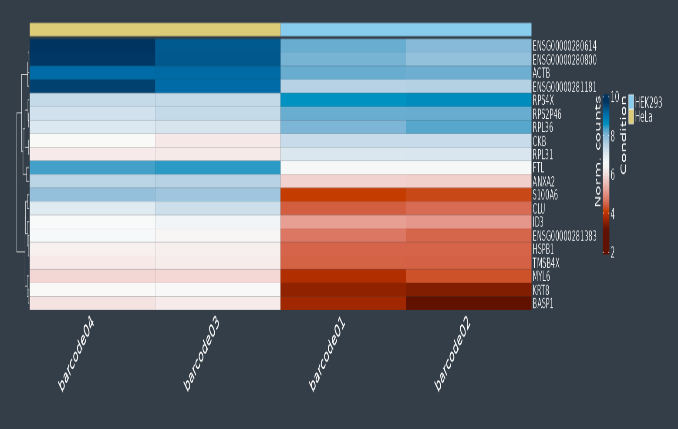

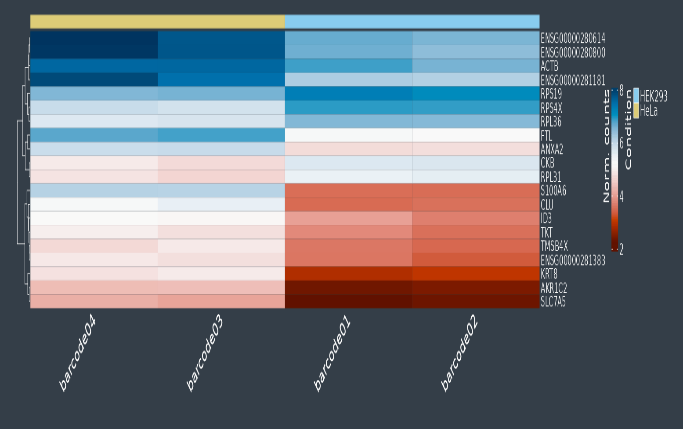

D

**Supplementary Figure S8- Comparison of “differential expression analysis” over time.** Iteration 2 (left) represents early sequencing state whereas iteration 60 (right) represents advanced sequencing state. **A.** Changes in Sample-2-sample variability throughout the sequencing. **B.** Changes in PCA plot throughout the sequencing **C.** Changes in DEGs represented in the Volcano Plot throughout the sequencing. **D.** Changes in top 20 differential expressed genes (based on p-adjusted) throughout the sequencing.

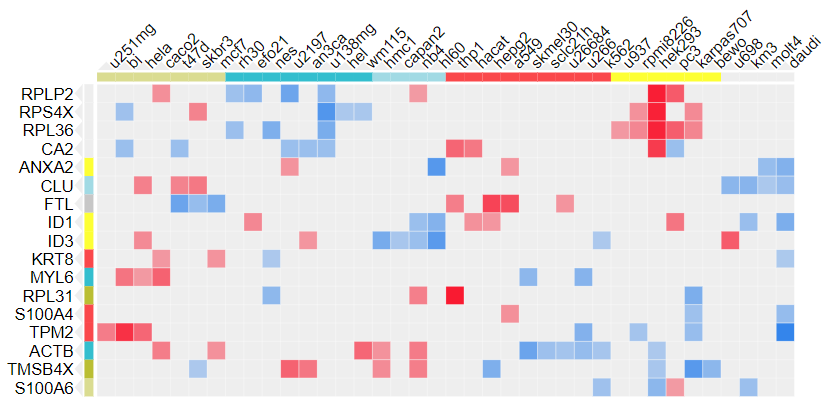

**Supplementary Figure S9- Confirmation of identified differentially expressed genes via database Harmonizome (Rouillard et al. 2016).** The Harmonizome cotains a collection of datasets including the “HPA Cell Line Gene Expression Profiles” that compares the differential gene expression across different cell line (Uhlén et al. 2015)**.** A list of the top 20 differentially expressed genes was compared to expression data of the Harmonizome database. The plot shows expression levels of the top 20 differentially expressed genes in different cell types including HEK293 and HeLa cells, which have been compared during the sequencing run. The red boxes represent gene enrichment within the cell line whereas blue boxes represent depletion. For additional information see Rouillard et al. 2016 and <https://maayanlab.cloud/Harmonizome/>
